# Phenotypic screening converges on CDK9 inhibition as a therapeutic strategy in translocation renal cell carcinoma

**DOI:** 10.1101/2025.08.25.672235

**Authors:** Dharma R. Thapa, Riva Deodhar, Adam Skepner, Prathyusha Konda, Meng Zhou, Joseph Bluck, Christian Pichlo, Yantong Cui, Paige Stocker, Prateek Khanna, Jiao Li, Jonathan M. Tsai, Alison Leed, Sandy Gould, Maria Alimova, Dhana Friedrich, Jaime H. Cheah, Rajesha Rupaimoole, Oivin Guicherit, Srinivas R. Viswanathan

## Abstract

Translocation renal cell carcinoma (tRCC) is an aggressive kidney cancer driven by gene fusions of the *TFE3* transcription factor. *TFE3* is essential in tRCC but dispensable in normal cells, presenting an attractive but pharmacologically challenging therapeutic target. We show that the basic helix-loop-helix (bHLH) domain of TFE3 is crucial for chromatin binding and transcriptional function. Via a phenotypic screen of 25,000 compounds, we identified molecules that either displace or retain chromatin-bound TFE3. BRD6866, a compound trapping TFE3 on chromatin, emerged as a pan-CDK inhibitor. Mechanistically, its inhibition of CDK9 – a key regulator of transcriptional elongation – was linked to impaired TFE3 fusion activity. These effects were recapitulated by the CDK9-selective inhibitor enitociclib, which downregulated TFE3 targets and suppressed tRCC cell growth. Our findings nominate CDK9 inhibition as a therapeutic strategy in tRCC and demonstrate the utility of mechanism-informed phenotypic screening for challenging targets.

## Introduction

Translocation renal cell carcinoma (tRCC) is a rare, aggressive, and molecularly-defined subtype of kidney cancer that comprises 1-5% of renal cell carcinomas (RCCs) in adults and 20-50% in children (1–3). There are no available systemic therapies that are specifically targeted to the biology of tRCC and patients with metastatic disease have poor survival (4,5) highlighting the urgent need for targeted therapies in this cancer.

Molecularly, the defining lesion in tRCC is an activating gene fusion involving an *MiT/TFE* family transcription factor gene (*TFE3*, *TFEB*, *TFEC*, or *MiTF*), with *TFE3* being rearranged in ∼90% of cases (5). Few other recurrent genomic alterations have been noted in tRCC, and the fusion therefore represents the dominant and often, sole, driver event (5–8). Oncogenic fusions in tRCC typically occur as in-frame fusions between *TFE3* and any one of >20 different partner genes (*ASPSCR1, NONO, SFPQ,* among many others), producing a chimeric fusion product (7,9–11).

TFE3 is a member of the MiT/TFE family of basic helix-loop-helix (bHLH) and leucine zipper (LZ) transcription factors (12), and in its wild type form, plays critical roles in regulating the transcription of genes involved in autophagy, lysosomal biogenesis and energy metabolism (13). Functionally, the bHLH domain mediates DNA binding to E-box consensus motifs at target gene promoters, while the LZ domain mediates homo/heterodimerization amongst MiT/TFE members (12–15). While many different partner genes and breakpoints have been identified in tRCC (6,7,10,16), the C-terminal region of TFE3, spanning exons 6-10 (which includes the bHLH/LZ domains), is uniformly retained in all fusions.

TFE3, like many other transcription factors, has limited structurally-resolved domains, no intrinsic enzymatic/ligand-binding activity, and no druggable pockets amenable to traditional structure-based or biochemical drug discovery approaches (17,18). For other similar transcription factors, recent studies have sought to overcome this liability by exploiting dependency on critical epigenetic regulators or transcriptional cofactors that may provide an alternative entry point to therapeutic targeting (19–22). For example, small molecule targeting and displacement of the bromodomain protein, BRD4, from chromatin is effective in models of androgen receptor (AR)-driven prostate cancer (23,24), *MLL/KMT2A* fusions driven leukemia (25,26), and in various MYC driven cancers (27). Inhibition of Menin, a chromatin associated protein, or DOT1L, a methyltransferase, has shown effectiveness in leukemia harboring *KMT2A* or *NUP98*-rearrangements (28–30). Likewise, targeting of transcriptional cyclin-dependent kinases (CDKs) (CDK7/CDK9/CDK11/CDK12/CDK13) has emerged as a therapeutic strategy in diverse cancers with dominant oncogenic drivers, including AR-driven prostate cancer (21) and various MYC-driven cancers (31–33).

Cell-based, phenotypic screening represents a powerful strategy to discover compounds that may disrupt transcription factor biology in a physiologic context via diverse mechanisms-of-action (MoA), including stability, sub-cellular localization, functional state, and protein-protein interactions (PPIs), amongst others. When combined with appropriate downstream biochemical and cellular assays for deconvolution, phenotypic screening can enable the discovery of both on-target and tightly-linked on-pathway hits and elucidate MoA for known inhibitors. For example, inhibition of Werner helicase (WRN), a well-validated target in microsatellite-unstable (MSI-H) cancers (34,35), appears to induce trapping of WRN on chromatin, leading to its SUMOlyation and subsequent ubiquitination and proteasomal degradation; phenotypic screens have elucidated the machinery regulating this process, which may offer alternative strategies for WRN targeting (36). Similarly, phenotypic characterization of the estrogen receptor (ER) inhibitor fulvestrant has revealed that its effects are not related merely to ER antagonism or degradation but to its ability to slow the intra-nuclear mobility of ER (37). These examples underscore the value of phenotypic screening as a versatile approach to uncover novel therapeutic strategies for challenging targets such as transcription factors – particularly those unamenable to enzymatic inhibition.

In this study, we leveraged a phenotypic screening strategy—previously applied in bromodomain inhibitor discovery (38) —to identify compounds capable of disrupting the DNA/chromatin binding activity of TFE3 fusion proteins. We find that small molecules can impair TFE3 fusion-driven transcription either by dislodging fusions from chromatin or by aberrantly trapping them on chromatin. Through this approach, we highlight CDK9, a transcriptional cyclin-dependent kinase (CDK), as a key therapeutic vulnerability in translocation renal cell carcinoma (tRCC).

## Results

### *TFE3* is selectively dependent in tRCC cells

The MiT/TFE fusion is a genetic hallmark of tRCC, with *TFE3* rearrangements being most common (6,10). Although many distinct *TFE3* fusion partners and breakpoint locations have been described (7,9,10), the resulting fusion transcripts typically retain at least Exons 6-10 of *TFE3*, which encode the basic helix-loop-helix (bHLH) and leucine zipper (LZ) domains essential for DNA binding and protein dimerization (14,39). Fusion products frequently also include coding sequences from the *TFE3* partner gene, which may incorporate additional functional domains (**Fig. 1A**).

**Figure 1.**
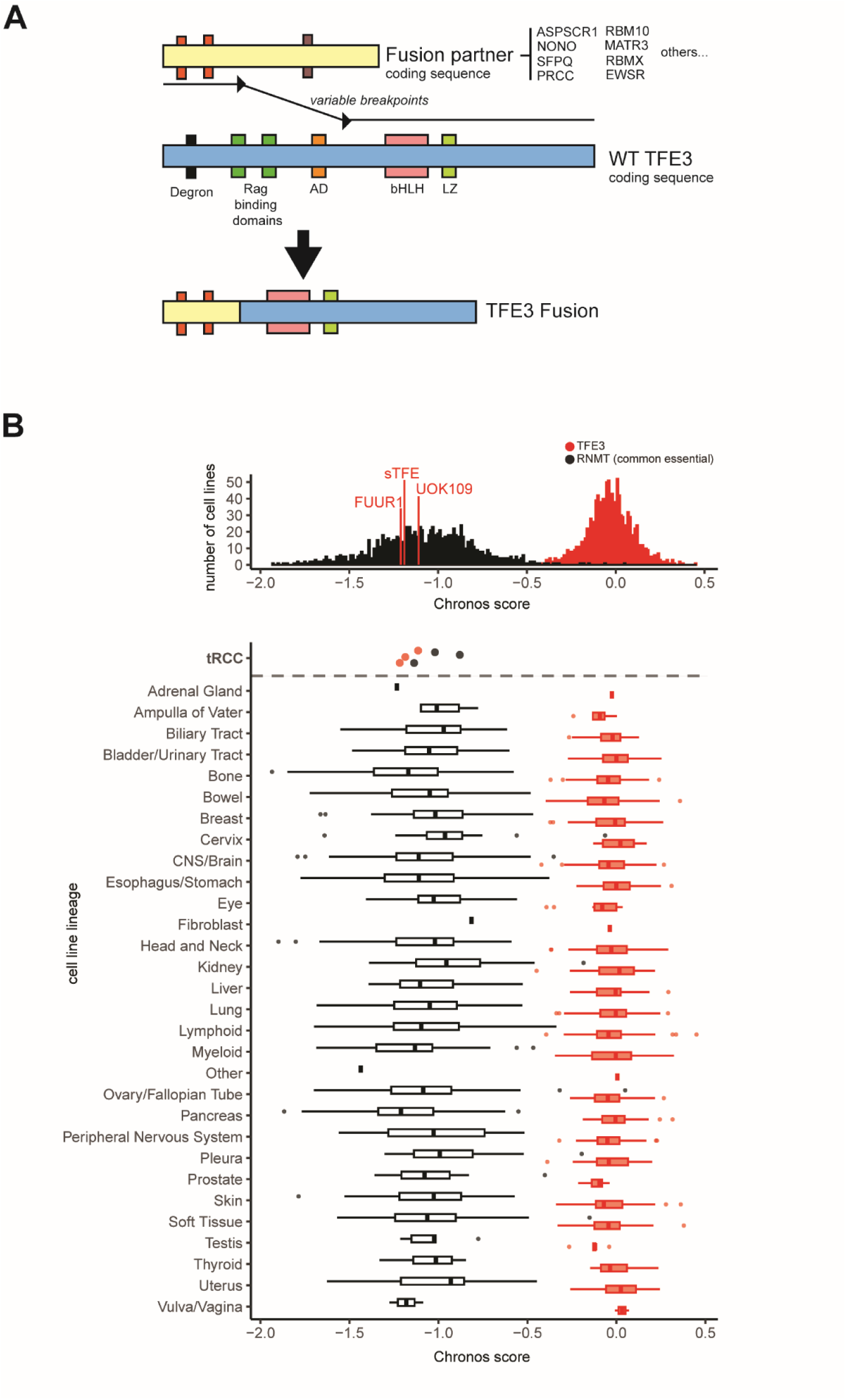
*TFE3* is a selective dependency in tRCC cells. **(A)** Schematic showing typical structure of *TFE3* fusions, which join coding regions of a variable fusion partner gene with the C-terminus of TFE3. Note: schematic shows retention of the minimal Exon 6-10 region of *TFE3* but upstream regions (e.g. including TFE3 activation domain, AD, and/or additional elements) are retained in some cases. **(B)** Chronos dependency scores for *TFE3* (red) and *RNMT* (representative pan-essential gene, black) across >1000 cell lines from 30 different lineages screened via genome-scale CRISPR screening in the Cancer DepMap (71). Chronos scores for *TFE3* and *RNMT* in three tRCC cell lines (FUUR1, STFE, UOK109) as separately determined by genome-scale CRISPR screening in a recent study (48) are shown in the bottom panel and highlighted in the histogram at top. Note: no tRCC cell lines are currently included in the Cancer DepMap.

The neomorphic structure of TFE3 fusions coupled with the normal phenotype of *Tfe3* knockout mice (40) raises the possibility of a favorable therapeutic window for the targeting of TFE3 fusions. We interrogated the Cancer Dependency Map (DepMap) dataset, which includes genome-scale CRISPR knockout screening data from >1000 cancer cell lines across many different lineages. Notably, although the DepMap includes more than 30 kidney cancer cell lines, tRCC cell lines have not yet been profiled (41,42). Gene dependency scores for *TFE3* were assessed using the Chronos metric (scaled to 0 for non-essential genes and −1 for essential genes) (43). In the DepMap, *TFE3* shows no strong dependency across all lineages, with scores distributed around 0, while *RNMT*, a representative pan-essential gene, has scores distributed around −1, confirming its essentiality (**Fig. 1B**). By contrast, genome-scale CRISPR screens conducted by our laboratory on three tRCC cell lines as part of a separate effort (44) reveal that *TFE3* exhibits strong dependency in these lines, similar to that of *RNMT*, with a mean Chronos score of −1.17. (**Fig. 1B**, red lines in histogram on top panel; first row of bottom panel). Thus, *TFE3* represents a strong, selective, and genetically-validated dependency in tRCC.

### The basic helix-loop-helix (bHLH) domain of TFE3 is critical for function

We sought to characterize the importance of regions within the bHLH/LZ domain, encoded within exons 7-9 and preserved in all oncogenic *TFE3* fusions. We generated several expression constructs for TFE3 or variants, including: TFE3 full-length (TFE3_FL), TFE3 spanning exons 6-10 (TFE3_Ex6-10), TFE3 with the bHLH region deleted (TFE3_ΔbHLH), and TFE3 with inactivating mutations in DNA-contacting residues (<u>n</u>on-<u>D</u>NA <u>b</u>inding, TFE3_NDB) (**Fig. 2A-B**). For the latter constructs, residues to be mutated were selected based on a homology model structure of the TFE3 dimer bound to a M-box DNA element, which was in the homology template structure MITF (PDB ID: 7D8T) (45). Six residues of TFE3 that display putative DNA interactions were mutated to alanine to generate TFE3_NDB (**Fig. 2B**).

**Figure 2.**
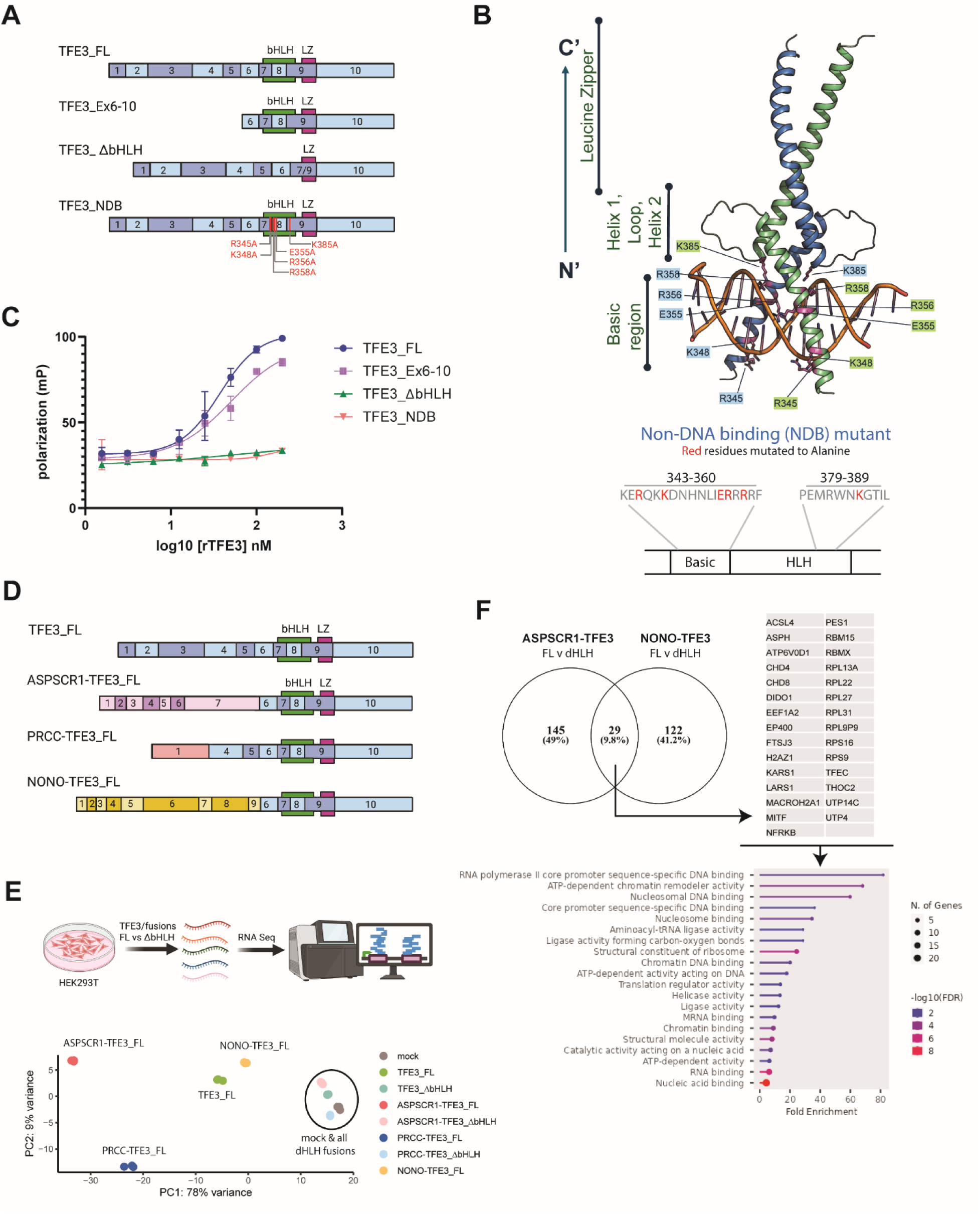
The basic helix-loop-helix (bHLH) domain of TFE3 is critical for DNA binding and downstream transcription. **(A)** Schematic of WT TFE3 and variants tested in this study. **(B)** Homology model of the bHLH and LZ domain of TFE3 (aa residues 343-432, based on the structure of MITF) with M-box DNA as a dimer. Purple residues represent residues within the bHLH domain that contact DNA, which were mutated in the non-DNA binding (NDB) mutant shown in (A). **(C)** Fluorescence polarization assay assessing binding of recombinant WT TFE3 or TFE3 variants to FAM-labeled E-box DNA element. **(D)** Schematic of full-length (FL) *TFE3* or various *TFE3* fusions tested in RNA-Seq study. The bHLH domain in all constructs were deleted to make the corresponding ΔbHLH variant. **(E)** Principal component analysis (PCA) from RNAseq performed after transient overexpression of the indicated constructs in HEK293T cells. RNA Seq schematic was created using BioRender.com. **(F)** Venn diagram and GO analysis of common interactors after BioID profiling of ASPSCR1-TFE3 and NONO-TFE3 fusions (FL vs ΔbHLH variant for each).

Purified recombinant TFE3 or its variants were tested for DNA-binding in a fluorescence polarization (FP) assay using FAM-labeled E-box DNA consensus sequence. TFE3_FL and TFE3_Ex6-10, which both have the bHLH domain, were capable of binding DNA, while TFE3_ΔbHLH and TFE3_NDB variants, with deletions or mutations in bHLH were deficient in DNA-binding (**Fig. 2C**). These findings confirm the necessity of the bHLH region in TFE3 for DNA binding and highlight the importance of specific amino acid residues in this interaction.

To determine if the bHLH region of TFE3 is essential for transcriptional activity, we generated full-length (FL) constructs for WT TFE3 and three different fusions (*ASPSCR1-TFE3*, *PRCC-TFE3*, and *NONO-TFE3*) (**Fig. 2D**), along with their corresponding ΔbHLH mutants. These were transiently expressed in HEK293T cells, followed by RNA extraction and RNAseq analysis. Principal component analysis (PCA) of RNAseq data showed that FL versions cluster separately from each other, while all ΔbHLH variants overlap with the mock condition, indicating a complete loss of their transcriptional activity (**Fig. 2E**). Validation with a doxycycline-inducible dominant negative variant of TFE3 (TFE3_DN) showed inhibition of TFE3 target gene activity in multiple tRCC cell lines within 8 hours of protein induction, genetically mimicking the effects of a potential TFE3 inhibitor (**Supplementary Fig. S1A**-**D**).

We then performed proximity interactome profiling (BioID) on ASPL-TFE3 and NONO-TFE3 fusions and their ΔbHLH variants. The interactome lost in the ΔbHLH variants of both fusions was enriched for nuclear DNA/RNA-binding proteins and transcriptional cofactors (**Fig. 2F**). These results are again consistent with the bHLH region of TFE3 fusions being crucial for DNA binding and transcriptional activity, providing a molecular rationale for targeting this function as a means to inhibit TFE3 fusions.

### Development of a phenotypic assay to detect TFE3 chromatin binding

While our studies implicate DNA-binding as the critical function to inhibit pharmacologically, using the biochemical FP assay employed above for screening could fail to capture entry points to inhibiting activity of the native TFE3 fusion complex on chromatin. Therefore, we turned to a cell-based, phenotypic approach that could assess the DNA-binding activity of TFE3 fusions, the chromatin displacement assay (CDA) (38). The CDA is a high-throughput, imaging-based method that has been previously employed for bromodomain drug discovery (38) (**Fig. 3A**); central to this protocol is the *in situ* cell extraction step, which removes proteins not bound to nuclear structures. Immunofluorescence and high-content imaging are used to quantify the nuclear signal of TFE3 with and without this harsh extraction. A decreased nuclear signal upon compound treatment (relative to DMSO) in the extraction condition indicates “displacement” of TFE3 from chromatin/DNA by the compound while an increased nuclear signal indicates its “retention.”

**Figure 3.**
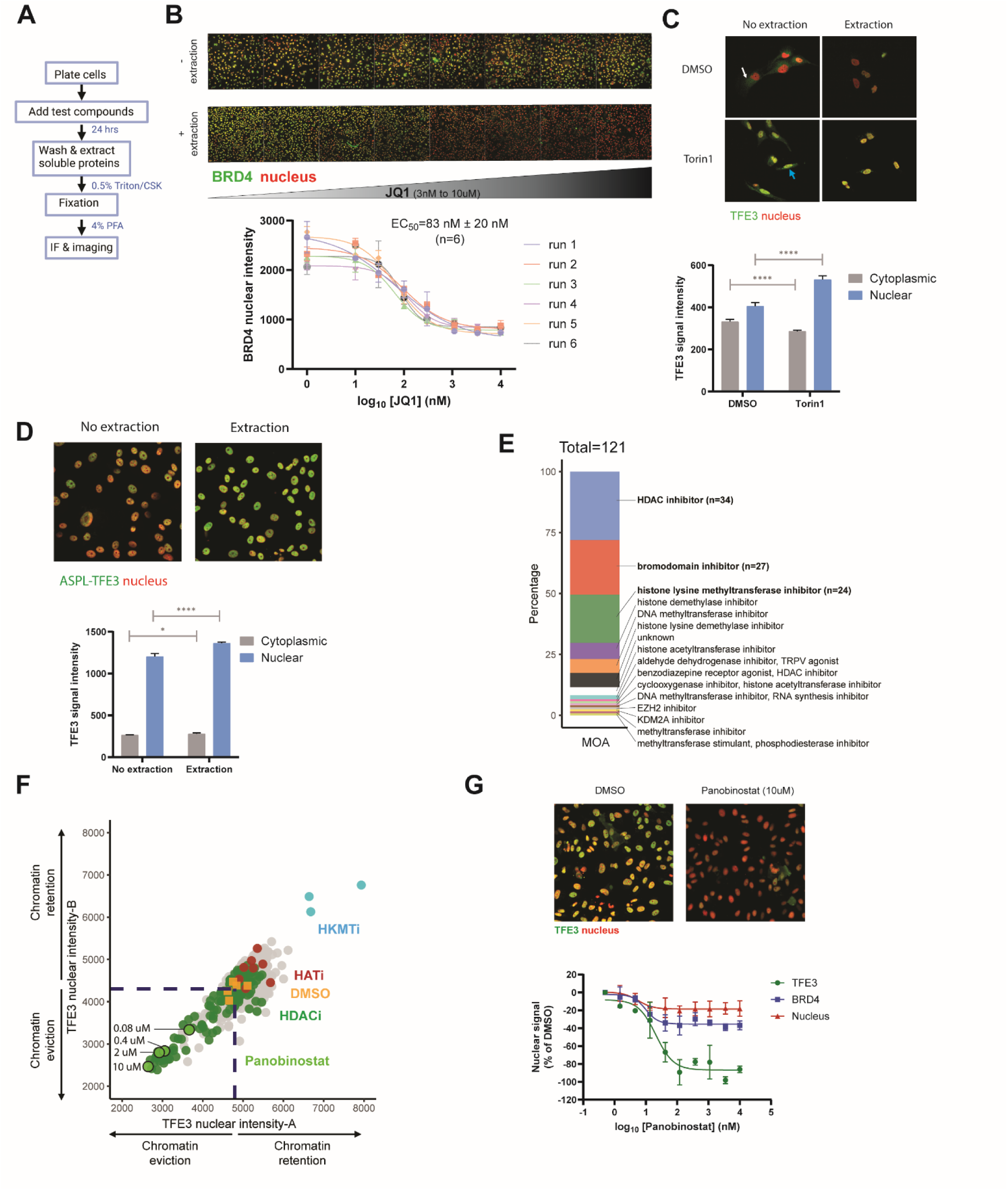
Adaptation of a cell-based chromatin displacement assay for phenotypic screening. **(A)** Outline of the chromatin displacement assay workflow. **(B)** Application of chromatin displacement assay to BRD4, using the tool compound JQ1. HEK293T cells were treated with JQ1 dose titration (3 nM to 10 μM). Cells were subject to *in situ* cell extraction (or not, bottom vs. top rows) and two color IF staining for BRD4 (green) and nucleus (red) was performed. Dose-dependent displacement of BRD4 from chromatin by JQ1 was quantified and plotted for each of 6 individual runs. Data are presented as mean ± SD. **(C)** Visualization of TFE3 staining and localization in HEK293T cells, with or without Torin1 treatment (which induces nuclear TFE3 localization), and with or without extraction. TFE3 cytoplasmic or nuclear intensity is quantified based on the IF images (mean ± SD, n=6). Note that cytoplasmic signal (white arrow) is dramatically reduced by extraction while nuclear (i.e. chromatin-bound) signal (blue arrow) is not. *P*-values computed by unpaired t-test; ****P<0.0001. **(D)** Visualization of ASPL-TFE3 fusion staining and localization in FUUR1 tRCC cells. Cytoplasmic and nuclear staining of the TFE3 fusion with or without extraction was quantified based on the IF images (mean ± SD, n=6) similar to panel (C). Note that TFE3 fusions are constitutively nuclear, as compared with WT TFE3, which shuttles between the cytoplasm and nucleus. *P*-values computed by unpaired t-test; *P<0.05, ****P<0.0001. **(E)** Composition of epigenetic modulator (“Epi-Mod”) compound library (n = 121 compounds) used for pilot screening in TFE3 chromatin displacement assay. **(F)** Replicate-Replicate scatterplot of Epi-Mod library screen using chromatin displacement assay in FUUR1 cells. Select inhibitor classes are highlighted: HDACi, green; HATi, red; HKMTi, blue. **(G)** Top, IF images showing displacement of ASPL-TFE3 fusion by panobinostat (10 μM) (with extraction condition). Staining: TFE3, green; Nucleus: Red. Quantification of dose-dependent displacement of TFE3, BRD4 and nuclear staining by panobinostat. Data is presented as mean ± SD, n=2.

As a first step toward adapting the CDA for TFE3, we first performed optimization using BRD4, a member of bromodomain and extraterminal (BET) family of proteins, which is known to be displaced from DNA by the specific tool inhibitor, JQ1 (38,46). In the extraction condition, JQ1 mediated a dose-dependent displacement of BRD4 from the nuclear signal across 6 independent runs (EC_50_ of 83 nM ± 20nM) (**Fig. 3B**).

To adapt this assay for TFE3, we used HEK293T cells to stain and detect WT TFE3 in both the cytoplasmic and nuclear compartment. Under nutrient replete conditions, WT TFE3 is labile (47) and has a strong cytoplasmic localization; treatment with the mTOR inhibitor, Torin1, mimics the cellular stress of nutrient starvation that leads to TFE3 stabilization and nuclear accumulation (**Fig. 3C**, left column). Under extraction conditions, TFE3 present in the cytoplasm is washed out, leading to a decrease in cytoplasmic signal. However, Torin1-activated nuclear localized TFE3 is retained in the nucleus even under extraction conditions suggesting tight nuclear binding of these TFE3 to chromatin (**Fig. 3C**, right column). In contrast to WT TFE3, TFE3 fusions are highly stable and predominantly nuclear. We validated the expression, stability, and nuclear enrichment of the ASPL-TFE3 fusion in FUUR1 tRCC cell line **(Fig. 3D)**. The nuclear expression of ASPL-TFE3 fusion is not disrupted by the extraction protocol confirming strong affinity of the fusion to nuclear and chromatin factors.

While JQ1 has emerged as a valuable tool compound to disrupt the chromatin-dependent activity of BRD4, such a tool compound does not currently exist for TFE3. We reasoned that, as a transcription factor, TFE3 might be dependent on certain epigenetic modifications for its chromatin localization. To identify an assay control for TFE3, we screened a small library of 121 diverse epigenetic modifiers (“Epi-Mod” library) (**Fig. 3E**) in the FUUR1 tRCC line. Each Epi-Mod compound was tested at 4 doses (10 μM, 2 μM, 0.4 μM, and 0.08 μM) using our CDA protocol (**Supplementary Table S1**). This revealed that histone deacetylase inhibitors (HDACi), particularly panobinostat, exhibited strong dose-dependent TFE3 fusion displacement from chromatin while their functional counterparts, the histone acetyltransferases inhibitors (HATi) showed increase in TFE3 chromatin signal (“chromatin retention”) **(Fig. 3F)**. We selected panobinostat for use as a displacement control in our CDA screening going forward (**Supplementary Fig. S2**).

In a follow up 10-point dose response study, panobinostat strongly displaced TFE3 fusion signal from the nucleus in a dose-dependent manner (**Fig. 3G)**. Intriguingly, despite not targeting TFE3 directly, panobinostat does show a degree of specificity for TFE3, and displayed much-reduced displacement of BRD4 (**Fig. 3G, green vs. blue**). Moreover, dose-dependent displacement of TFE3 fusion from chromatin was accompanied by a minimal decrease in cell viability based on change in nuclei count over the 24 hour time course of the experiment (**Fig. 3G, red**). Thus, the CDA can be adapted to the oncogenic TFE3 fusion and we identified panobinostat as an assay positive control for TFE3 displacement.

### 25,000 compound screen for inhibitors of TFE3 fusion activity

Having established the CDA for TFE3 fusions, we proceeded with a screen of a diverse compound library of ∼25,000 molecules (in-house curated diversity set of molecules with drug-like properties), in order to identify additional hits that can displace the ASPL-TFE3 fusions from chromatin in FUUR1 cells (**Fig. 4A**). Each compound was screened at single dose (10 μM) in duplicate and the entire library was screened across 5 separate batches with overall robust assay performance (aggregate signal/noise (S/N) ratio range of 1.5-1.8; Z’ factor range of 0.2-0.7). Hits were called for each batch using a cutoff for displacement if values <DMSO (median - 3*RSD) and retention if values >DMSO (median + 8*RSD) which resulted in cumulative 62 hits from the primary screen (**Supplementary Fig. S3A, Supplementary Table S2**). Of these, 44/62 (71%) led to decreased TFE3 nuclear signal vs. DMSO with extraction (“chromatin displacement”) while 18/62 (29%) led to increased TFE3 nuclear signal vs. DMSO with extraction (“chromatin retention”) **(Fig. 4B, red and green dots)**. After 8-point dose response validation, 4/62 primary hits (6%) showed dose-dependent effect on TFE3 nuclear signal. Of these, 3 hits scored in the chromatin displacement mode (BRD7659, BRD7186, BRD4539) while the remaining hit (BRD6866) scored in the chromatin retention mode (**Supplementary Fig. S3A**). Notably, while the three chromatin displacement hits were active only at top dose, the retention hit, BRD6866, showed a pronounced dose-dependent chromatin trapping of TFE3 (**Fig. 4C-E** and **Supplementary Fig. S3B**). Another outlier hit in the retention mode in the primary screen, BRD5169, did not validate in 8-point dose response validation (**Supplementary Fig. S3B**). We also tested 5 additional compounds in dose response based on their ability to selectively kill tRCC cells vs. other kidney cancer cells in a separate drug repurposing library screen (“Repo screen”) (48) (**Supplementary Fig. S3C-D**). Of these, only bardoxolone, previously characterized as an activator of the NRF2 pathway (49), validated in dose-response, displaying a dose-dependent TFE3 chromatin retention phenotype similar to BRD6866.

**Figure 4.**
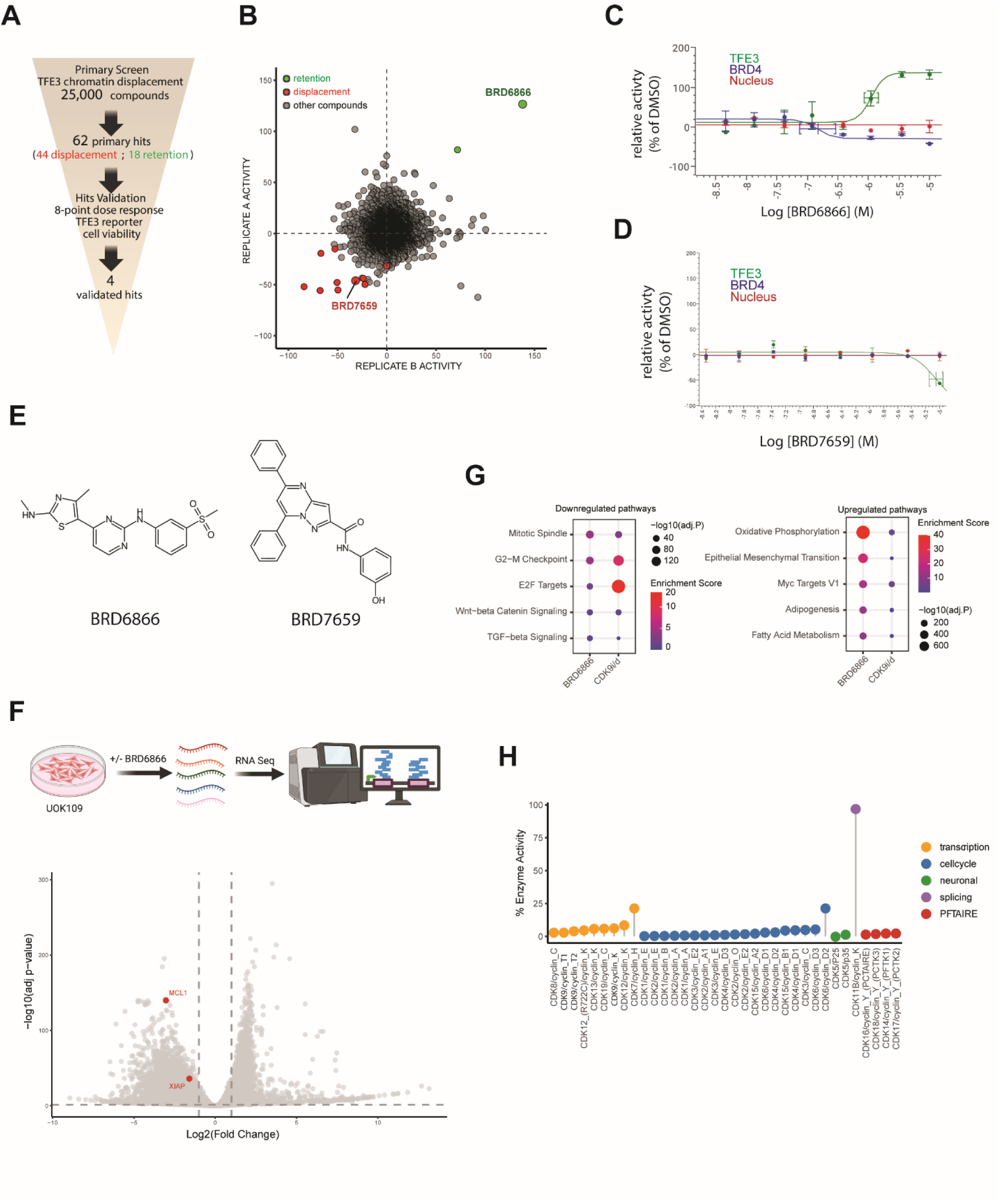
25,000 compound diversity screen in tRCC cells using the TFE3 chromatin displacement assay. **(A)** Screening funnel of 25,000 compounds leading to 4 hits after dose response validation studies. **(B)** Replicate-Replicate scatterplot for run 4 (out of 5 total runs) of chromatin displacement screening of 25,000 compounds in in FUUR1 cells. Chromatin displacers, red; chromatin retention, green (see methods for hit selection from the 5 separate runs). Primary hits from this run are labeled in red or green; validated hits are named. **(C)** Dose-dependent increase in nuclear signal of TFE3 with BRD6866 treatment (chromatin retention) in a chromatin displacement assay (CDA) in FUUR1 cells. Nuclear signal (DAPI) as well as nuclear IF signal for TFE3 and BRD4 were quantified and plotted (mean ± SD, n=3). **(D)** Dose-dependent decrease in nuclear signal of TFE3 with BRD7659 (chromatin displacement) in a CDA in FUUR1 cells. Nuclear signal (DAPI) as well as nuclear IF signal for TFE3 and BRD4 were quantified and plotted (mean ± SD, n=3). **(E)** Chemical structure of BRD6866 and BRD7659. **(F)** RNA Seq of UOK109 cells after 16h treatment with BRD6866 at 1μM. Differential transcriptomics analysis using a volcano plot is shown. Antiapoptotic genes with known sensitivity to CDK9 inhibition, MCL1 and XIAP are labeled (51). RNA Seq schematic created using BioRender.com. **(G)** Hallmark gene set pathway enrichment analysis upon BRD6866 treatment was compared to two recent studies with CDK9 inhibition (CDK9i) and CDK9 degradation (CDK9d) (21,32). **(H)** CDK activity profiling assay (Reaction Biology) showing extent of inhibition of various CDK/cyclin pairs by BRD6866 (10 μM). Loss of activity for each CDK and cyclin pairs by BRD6866 was compared to DMSO control. CDK9/cyclin pairs are bolded. The cyclin interacting motif PFTAIRE defines a subgroup of CDKs that does not fall under named categories.

BRD6866 is an aminopyrimidine molecule (**Fig. 4E**), a structural motif previously linked to inhibition of transcriptional CDKs (50), and which had been annotated within our library as an inhibitor of multiple cyclin dependent kinases, including CDK2 and CDK9. To further characterize the mechanism of action of this compound, we performed whole transcriptomic analysis after treatment of UOK109 cells with BRD6866 (**Fig. 4F**). The antiapoptotic transcripts from the BCL2 family -- *MCL1* and *XIAP*, both known to be sensitive to CDK9 inhibition (51) -- were strongly downregulated by BRD6866. Furthermore, comparison of Hallmark pathway enrichment analysis from BRD6866 treated cells with CDK9 inhibitior (21) or CDK9 degrader (32) treated cells in prior studies showed similar pathways downregulated in all conditions (mitotic spindle, G2-M checkpoint, E2F targets) or upregulated (oxidative phosphorylation, Myc targets, epithelial mesenchymal transition) suggesting a strong CDK9 inhibitory property of BRD6866 (**Fig. 4G**). We then performed a cyclin-dependent kinase (CDK) profiling assay of BRD6866 against a panel of 35 different CDKs and cyclin pairs. The profiling revealed BRD6866 to be a potent pan-CDK inhibitor across all families of CDKs, including both cell-cycle and transcriptional CDKs (the latter of which regulate RNA polymerase II (RNAP II)-mediated transcriptional elongation and mRNA maturation (52)) (**Fig. 4H**).

### CDK9 inhibition suppresses TFE3 activity and induces cell death of tRCC lines

To investigate whether altering TFE3 fusion chromatin dynamics (either via displacement or retention) was associated with a decrease in target gene expression, we constructed a dual luciferase reporter in UOK109 cells, with Firefly luciferase driven by a TFE3-responsive element (promoter region from the *TFE3* target gene, *GPNMB*) (53) and Renilla luciferase driven by an *EF-1*α promoter (non TFE3-responsive control).

While BRD7659, BRD7186, BRD4539 had minimal effects on TFE3-reporter activity (**Supplementary Fig. S4A**), BRD6866 led to a dose-dependent decrease in TFE3 reporter activity (**Fig. 5A**). Intriguingly, this was despite its <u>increasing</u> TFE3 chromatin signal in a dose-dependent fashion, suggesting that it may act by “trapping” TFE3 fusions on chromatin and impairing their downstream transcriptional activity. Moreover, the IC50 for suppression of the TFE3-responsive reporter was ∼4-fold lower than that for the control reporter (IC_50_ of 342 nM vs 1388 nM), suggesting selectivity for repression of TFE3 target genes versus a general repression of transcription.

**Figure 5.**
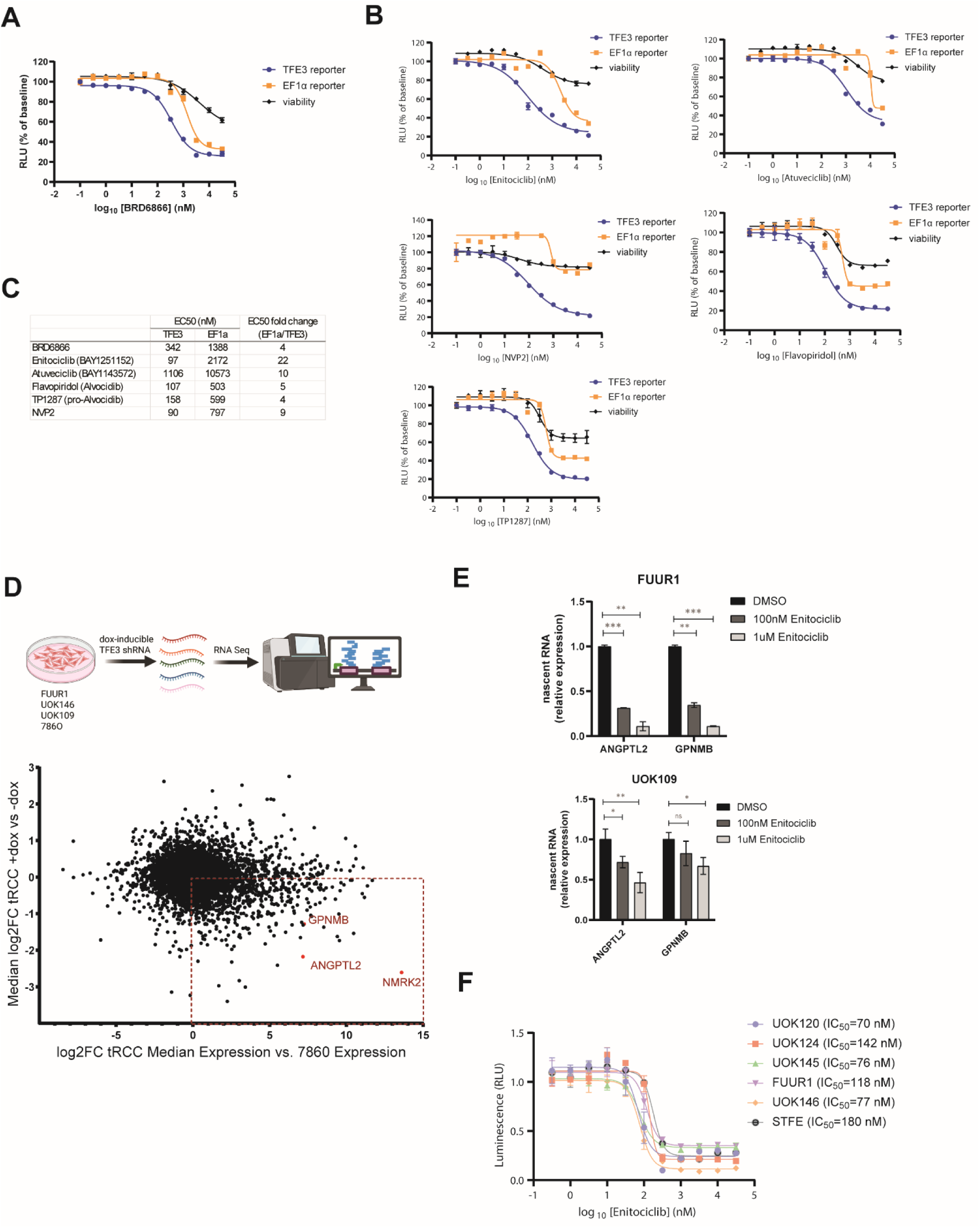
Mechanistic validation of hits from primary screen implicates CDK9 inhibition as strategy to suppress TFE3 fusion activity in tRCC. **(A)** Dual reporter luciferase assay reveals selective inhibition of TFE3 fusion-driven reporter activity (blue) vs. control reporter activity (EF1α, yellow) by BRD6866. Viability (CellTiterGlo) is shown in black. Reporter assay performed in the UOK109 tRCC cell line expressing the NONO-TFE3 fusion after 24 hrs of drug treatment (mean ± SD, n=3). **(B)** Dual reporter luciferase assay in UOK109 cells with a panel of well-characterized CDK9 inhibitors (mean ± SD, n=2). The legend color coding for TFE3 and EF1α reporter activity and viability is identical to A. **(C)** EC50 values for activity suppression with TFE3 and EF1α reporters for the compounds in A and B. **(D)** RNA Seq analysis of tRCC cells (FUUR1, UOK146, UOK109) and ccRCC cell (786O) 7 days after induction of TFE3 shRNA by doxycycline. Dotted area in scatter plot shows gene transcripts that are highly expressed and responsive (downregulated) to TFE3 knockdown in tRCC lines compared to 786O. RNA Seq schematic created using BioRender.com. **(E)** Nascent RNA of canonical TFE3 fusion target genes (*ANGPTL2*, *GPNMB*) was measured by RT-qPCR after 2 hours of enitociclib treatment in FUUR1 (left) and UOK109 (right) cells (mean ± SD, n=3). *P*-values computed by unpaired t-test; *P<0.05,**P<0.01,***P<0.001. **(F)** Cell viability assay in various tRCC cell lines treated with the indicated doses of enitociclib for 72 hrs (mean ± SD, n=2)

To deconvolute the relative contribution of cell-cycle vs transcriptional CDK inhibitory activity of BRD6866 on TFE3 reporter inhibition, we tested several cell cycle active CDK4/6 inhibitors in our dual reporter assay: Palbociclib (CDK4/6), Abemaciclib (CDK4/6), and PF-07220060 (CDK4). All showed minimal inhibition of TFE3 reporter activity while the transcription regulating CDK inhibitor THZ-531 (CDK12/13) showed modest inhibition of TFE3 reporter activity (**Supplementary Fig. S4B**). This finding, coupled with recent studies linking CDK9 inhibition to oncogenic transcription (54,55), prompted to investigate CDK9 inhibition as the primary mechanism of TFE3 fusion target gene suppression.

We therefore next tested multiple highly selective CDK9 inhibitors in our dual reporter assay (enitociclib, atuveciclib, NVP2, flavopiridol, and TP1287), finding that all showed dose-dependent selective inhibition of the TFE3-responsive reporter compared to the EF-1α reporter in our dual reporter assay (**Fig. 5B**). This difference was most pronounced with enitociclib (VIP152), a potent and selective inhibitor of CDK9 currently in clinical trials (NCT05371054) (56), which showed >20x activity against the *TFE3* reporter vs. the control reporter (**Fig. 5C**). In addition, we observed that multiple of these CDK9 inhibitors showed a dose-dependent TFE3 displacement phenotype in the chromatin displacement assay, indicating that CDK9 inhibition can be tied to altering TFE3 fusion dynamics on chromatin (either via displacement or via retention, as with BRD6866) (**Supplementary Fig. S4C**).

Given the role for CDK9 in regulating early transcription (e.g. transition from transcriptional initiation to elongation), we wanted to assess nascent RNA transcripts from TFE3 target genes by qPCR. TFE3 target genes for evaluation were selected based on those that were highly expressed in tRCC lines (FUUR1, UOK146, UOK109) and responsive to shRNA mediated knockdown of *TFE3* when compared to a ccRCC line (786–0) (**Fig. 5D**). Enitociclib treatment in both FUUR1 and UOK109 cells resulted in a dose-dependent decrease in the levels of nascent transcripts for two of these canonical TFE3 target genes (*ANGPTL2*, *GPNMB*) (**Fig. 5E**). We next treated a panel of tRCC lines harboring various *TFE3* fusions (FUUR1 and STFE: *ASPSCR1-TFE3*; UOK120, UOK124, and UOK146: *PRCC-TFE3*; UOK145: *SFPQ-TFE3*) with enitociclib *in vitro*. After 72 hours of treatment, all tRCC lines showed high sensitivity to CDK9 inhibition in the low nM range (IC_50_ range: 70.32 nM to 179.8 nM), comparable to other sensitive cell lineages such as lymphoma (57) (**Fig. 5F**).

## Discussion

In this study, we present a mechanism-driven high-throughput screening strategy to identify inhibitors of TFE3 fusion activity. The TFE3 fusion represents a high-value target with a wide therapeutic index in a genetically-defined cancer (tRCC) with no effective therapies (10,58–60). We validate a requirement for the bHLH/LZ domain in DNA binding and transcriptional activity of TFE3 fusions and used this finding to inform development of a novel cell-based, phenotypic screening approach to identify molecules that can modulate the DNA/chromatin binding of TFE3 fusions in its native setting.

This approach identified CDK9 as a potential therapeutic target in tRCC. CDK9 in association with cyclin T1 constitutes the catalytic subunit of the positive transcription elongation factor b complex (p-TEFb), which phosphorylates the RNA polymerase II C-terminal domains (CTD) at Ser2 and promotes transcriptional elongation of most genes (61,62). While p-TEFb is universally required for transcription, some cancer types, particularly those dependent on strong oncogenic drivers or on the elevated expression of short-lived transcripts such as *MCL1* and *MYC* (22), are particularly vulnerable to CDK9 inhibition (63,64). Indeed, inhibition of CDK9 has shown potent antitumor efficacy in multiple cancer types including chronic lymphocytic leukemia (CLL) (65), MYC-driven lymphoma (66), MYC/MCL-1-driven multiple myeloma (67), and AR-driven prostate cancer (21), amongst others (68). In this study, we extend this observation of CDK9 dependence to tRCC, where the TFE3 fusion represents a strong oncogenic transcription factor driver analogous to AR or MYC in other CDK9-dependent cancer types; whether specific TFE3 fusion target genes are chiefly responsible for CDK9-dependence represents an exciting avenue for future investigation.

Surprisingly, a top hit from our screen, BRD6866 scored not by displacing TFE3 from chromatin, but by inducing its retention; other, more selective, CDK9 inhibitors also scored in our assay, but in a displacement mode. The mechanism by which CDK9 inhibition alters the dynamics of TFE3 on chromatin remains to be defined. Nonetheless, these data add to a growing body of evidence that trapping nuclear proteins on chromatin can impair their function, and this mechanism may be as or more effective than enzymatic inhibition or degradation alone: clinically-relevant examples include WRN inhibitors, fulvestrant, and some PARP inhibitors (37,49,69). Moreover, our screen suggests that trapping compounds can potentially be discovered with higher robustness than those inducing displacement, perhaps because chromatin retention represents a positive-selection, or “up” assay (70). Assay performance suggests that this high-throughput screening approach would be amenable to the screening of larger and more diverse chemical libraries in future studies.

Our screening strategy implicates CDK9 as a required cofactor for TFE3 fusion transcriptional activity and suggests CDK9 inhibition as a rational therapeutic strategy in tRCC. Although direct inhibitors of TFE3 were not identified in our study, screening of a larger library with an appropriate deconvolution cascade may identify such molecules. More broadly, we present a phenotypic screening approach that may find utility in the discovery of inhibitors for other challenging transcription factor targets.

## Methods

### Cell lines and plasmids

FUUR1 was obtained from Dr. Masako Ishiguro’s laboratory (Fukuoka University School of Medicine), UOK109, UOK146, UOK120, and UOK124 from Dr. W. Marston Linehan’s laboratory (National Cancer Institute), STFE from RIKEN (# RCB4699), and HEK293T from ATCC® (CRL-11268). All cell lines were cultured at 37°C in DMEM “complete media” with 10% FBS, 100 U/mL penicillin, and 100 μg/mL Normocin (Thermo Fisher: #NC9390718).

Full length and ΔbHLH versions of TFE3, ASPL-TFE3, PRCC-TFE3, NONO-TFE3 were initially cloned into the Gateway system using pDONR221 (ThermoFisher #12536017) to create Entry vectors. For overexpression studies, the entry clones were transferred into the destination vector pLX_313 (Broad Institute) using Gateway system (ThermoFisher #11791100). For BioID studies, the BirA* (BirA R118G) open reading frame was first cloned downstream of the attR2 sequence in the pLX_313 vector to create the pLX_313_BirA* destination vector. All entry clones were then transferred to pLX_313_BirA* destination vector for use in the BioID study. TFE3_DN was synthesized into pTwistENTR vector (Twist Biosciences) and cloned into pInducer20 (Addgene #44012) using the Gateway system (ThermoFisher #11791100). For shRNA studies, TFE3 targeting shRNA (sequence: AATCAGATAAACAAATGAGGG) was cloned into doxycycline-inducible lentiviral vector as previously described (73).

### Transfection, lentiviral production and stable lines

All transient transfections were performed using Lipofectamine^TM^ 2000 (ThermoFisher #11668019). For generation of stable lines, lentivirus was first prepared by transfecting HEK293T cells with pInducer20_TFE3_DN and packaging plasmids (psPAX2, pMD2.G) using polyethylenimine-based transfection. Transfection media was replaced 14-16 hours later with complete growth media and the supernatant containing the virus was harvested 3 days post-transfection. Virus was stored at - 80°C until use. tRCC cell lines (FUUR1, UOK146, UOK109) or ccRCC line (786O) were infected with viral supernatant using a spinfection method. Briefly, virus was added to the cells, mixed with polybrene (1.5 ug/ml) and infected by centrifugation for 800g at 33°C for 1.5 hours. The cells were then washed with media and cultured for 2-3 days before starting selection using Hygromycin (100 μg/ml) (ThermoFisher #10687010) or G418 (500 μg/ml) (ThermoFisher #10131035) until stable clones were generated.

### Fluorescence polarization assay

Recombinant TFE3 variants bacterial expression vectors were custom ordered from GenScript. In brief, the proteins were then overexpressed in E. coli in Luria Broth medium and purified from the supernatant of cell lysates using Ni column/Chromdex 200 columns and resuspended in storage buffer (50 mM Tris-HCl, 500 mM NaCl, 10% Glycerol, pH 8.0). Varying concentration of the recombinant TFE3 variants were mixed with 5 nM of FAM-labeled Ebox DNA (CCAT**CACATG**ATCCT) in black opaque non-binding flat bottomed 96 well plate (Greiner, #655900) and incubated for 3 hours at 4°C in binding buffer (20 mM HEPES, 200 mM NaCl, 1 mM MgCl_2_, 2.5% glycerol). The binding between the TFE3 variants and FAM-labeled Ebox DNA was measured by fluorescence polarization using CLARIOstar^plus^ microplate reader. Polarization (P) was calculated from emission fluorescence intensity (F) that is parallel and perpendicular to excitation light plane using the equation: P = (F_parallel_ -F_perpendicular_)/(F_parallel_ + F_perpendicular_) and expressed as millipolarization (mP) where mP = 1000 P.

### Chromatin displacement assay and immunofluorescence

384-well plates (Perkin Elmer PhenoVue) were coated with Synthemax II-SC substrate (Corning #3535). Test compounds were diluted to 10 mM stock and 40 nL was echo transferred to wells to create assay ready plates (ARPs). HEK293T or FUUR1 cells (7,000 cells/well) were seeded into ARPs in 40 μL of media (reconstituting compounds to final concentration of 10 μM) and cultured overnight at 37°C/5% CO_2_ incubator. 24 hours later, the plates were washed with PBS (no Mg^2+^/Ca^2+^) followed by freshly prepared ice-cold cytoskeleton (CSK) buffer (10 mM PIPES, 300 mM sucrose, 100 mM NaCl, 3 mM MgCl_2_; pH = 6.8). In the next step, extraction of non-chromatin-bound proteins, was performed by addition 50 μL of cold extraction buffer (CSK buffer + 0.8% Triton X-100) for 10 min at 4°C. The cells were then fixed for 20 minutes at room temperature by addition of 16% paraformaldehyde (PFA) directly to the extraction buffer to obtain 4% PFA. A second fixation step was performed with 4% PFA for an additional 10 minutes at room temperature followed by three washes with PBS (no Mg^2+^/Ca^2+^).

For immunofluorescence staining, the fixed cells were blocked (PBS + 1% BSA) for 30 minutes at room temperature and stained with primary antibodies for 1 hour at 37°C at the following dilutions: rabbit anti-human TFE3 antibody (Millipore Sigma #ZRB1272) at 1:1000 and mouse anti-human BRD4 antibody (Santa Cruz Biotechnology #sc518021) at 1:200. After 3x washing with PBS (no Mg^2+^/Ca^2+^), the cells were incubated in secondary antibody for 1 hour at room temperature in the dark using the following dilutions: goat anti-rabbit IgG (H+L) Alexa Fluor^TM^ Plus 488 (ThermoFisher #A32731) at 1:1000; donkey anti-mouse IgG (H+L) Alexa Fluor^TM^ 647 (ThermoFisher #A31571) at 1:1000; Hoescht 3342 (ThermoFisher #H3570) at 1:1000. The plates were then washed 3x with PBS (no Mg^2+^/Ca^2+^) and sealed with the wash buffer for storage at 4°C until imaging with Perkin Elmer Pheonix Opera instrument with settings—Objective:20X; Fields/well: 4 to 9; channels: TFE3-AF488, BRD4-AF647, Hoechst, and Brightfield. The 25,000 library was covered by 158 x 384 well plates which was run in 5 batches with robust assay statistics (Z’ factor between 0.2-0.7; signal/noise (S/N) ratio between 1.5-1.8). Displacement hits were called if mean between replicas were < DMSO controls (median – 3 * RSD) and retention hits if mean between replicas > DMSO controls (median + 8 * RSD). Validation runs were performed using 8-point, 3-fold dose response starting at 25 μM using the exact protocol used in the screen.

### Western Blot

Cell lysis was performed using ice-cold RIPA buffer (ThermoFisher, #89901) supplemented with protease inhibitor (Roche, #11836170001) and phosphatase inhibitors (Roche, #4906845001). The lysate was probe sonicated for 3 x 30 second bursts on ice to ensure complete disruption of cellular contents and centrifuged at 14,000 x g for 15 minutes to obtain the total soluble protein which was then quantified using the Pierce BCA protein assay kit (ThermoFisher, #23225). Equal amounts of protein were loaded onto Bolt 4-12%, Bis-Tris Plus WedgeWell gels (ThermoFisher, NW04125BOX) for SDS-PAGE separation at 150V for 1h. Proteins were transferred to a nitrocellulose membrane using the iBlot 2 dry transfer method (ThermoFisher). The membrane was blocked (LI-COR, #927-60001) for 1 hour and incubated (with gentle rocking) overnight at 4 °C with the primary anti-HA tag antibody (Cell Signaling Technology, #3724S) or anti-b-actin antibody (Cell Signaling Technology, #8457L) in antibody diluent buffer (LI-COR, #927-65003). The following day, the blots were washed (3x with TBS-T) and incubated for 1 hour with secondary goat-anti-rabbit-HRP (Abcam #ab6721) in antibody diluent buffer at room temperature. After three TBS-T washes, the blot was incubated with the Pierce ECL substrate (ThermoFisher #32209) for signal generation and was imaged using the Odyssey M Chemiluminescent Imaging System (LI-COR Biosciences).

### Metabolic labeling of nascent RNA

Nascent gene-specific mRNA synthesis was measured by using the Click-iT Nascent RNA Capture Kit (ThermoFisher, #C10365). FUUR1 or UOK109 cells were treated with 100 nM or 1000 nM of enitociclib (MedChem Express, #HY-103019E) followed by the addition of 5-ethyly uridine (EU) for 2 hours. The EU labeled RNA was collected with the Quick RNA mini-prep kit (Zymo Research, #R1055) and biotinylated in a copper-catalyzed click reaction followed by nascent RNA capture with streptavidin magnetic beads based on recommended protocol. The bead-captured EU nascent RNA was reverse transcribed to cDNA using Superscript IV VILO Master Mix (ThermoFisher, #11756050) and analyzed by RT-qPCR.

### Real-time quantitative qPCR (RT-qPCR) and RNA-seq

Total RNA was extracted using the Quick RNA mini prep kit (Zymo Research, #R1055) followed by cDNA synthesis from 1ug of total RNA using superscript IV VILO Master Mix (ThermoFisher Scientific, #11756050). The qPCR reactions were performed using 10ng of cDNA with the TaqMan Fast Advanced Master Mix (Thermo Fisher Scientific #4444557) and the following gene-specific primers (ThermoFisher, #4331182): *GAPDH* (#Hs02786624_g1), *GPNMB* (#Hs01095669_m1), *ANGPTL2* (#Hs00171912_m1), *PKD1L2* (#Hs00385217_m1), *NMRK2* (Hs01043681_m1), *TRIM63* (Hs00822397_m1). Reactions were run on the StepOne Real-Time PCR System (Applied Biosystems) according to the recommended protocol. Expression levels of each gene was normalized to GAPDH and fold change was calculated relative to DMSO treated samples using the ΔΔCt method. For RNA-seq, total RNA was sent to Novogene where the samples were sequenced using the NovaSeq X Plus Series (PE150) platform and strategy. Paired-end sequencing reads were aligned to the human genome reference (GRCh38.p13 assembly) using STAR (v2.7.10a) and quantified as paired-end reads against the GENCODE v45 transcript reference using RSEM (v1.3.1). Transcripts were filtered based on read support (sum of read counts across three biological replicates > 30) prior to gene-level differential expression analysis using DESeq2 (v1.34.0) using default parameters (74). Thresholds for significant down/upregulation were defined as q < 0.05 and |log2fold-change| > 1. The counts were normalized via variant stabilizing transformation (vst) and principal components were calculated to visualize the effect of experimental covariates using plotPCA function.

### Luciferase reporter assays

UOK109 line with stable expression of GPNMB (a TFE3 target) promoter-driven NanoLuc was infected with lentivirus expressing EF1α-driven Renilla luciferase (pLX313-RLuc, Addgene #118016) to create a dual reporter line. 10,000 cells in 100uL of DMEM complete media were seeded per well in a 96 well white plate (Corning, #3903). The following day, compounds or DMSO were added to the indicated concentration. After 24 hours of treatment, signals for NanoLuc (Nano-Glo system, Promega, #N1110), Renilla Luciferase (Renilla-Glo system, Promega, #E2710), and cell viability (CellTiter-Glo assay, Promega, G7570) were quantified using the CLARIOstar^plus^ microplate reader (BMG labtech) following manufacture’s protocol.

### *In vitro* kinase activity assay

Testing for inhibition of CDK activity was performed by Reaction Biology Corporation using their Kinase HotSpot Assay. BRD6866 was tested at a single dose (10 uM) against a panel of 34 CDK/cyclin pairs. Staurosporine was used as control and tested in 10-dose with 4-fold serial dilution starting at 20 uM. AT-7519 and THZ531 was used as alternate controls for CDK11B/cyclin K and CDK12-13/cyclin K respectively. All reactions were carried out at Km ATP for the CDK/cyclin pairs. Cyclin1/cyclin A (10 μM ATP), CDK1/cyclin B (30 μM ATP), CDK1/cyclin E (30 μM ATP), CDK2/cyclin A1 (30 μM ATP), CDK2/cyclin A (20 μM ATP), CDK2/cyclin E (30 μM ATP), CDK2/cyclin E2 (100 μM ATP), CDK2/cyclin O (50 μM ATP), CDK3/cyclin C (5 μM ATP), CDK3/cyclin E (50 μM ATP), CDK3/cyclin E2 (50 μM ATP), CDK4/cyclin D1 (100 μM ATP), CDK4/cyclin D2 (100 μM ATP), CDK4/cyclin D3 (30 μM ATP), CDK5/P25 (15 μM ATP), CDK5/p35 (50 μM ATP), CDK6/cyclin D1 (100 μM ATP), CDK6/cyclin D2 (100 μM ATP), CDK6/cyclin D3 (100 μM ATP), CDK7/cyclin H (50 μM ATP), CDK8/cyclin C (20 μM ATP), CDK9/cyclin K (15 μM ATP), CDK9/cyclin T1 (10 μM ATP), CDK9/cyclin T2 (30 μM ATP), CDK11B/cyclin T2 (30 μM ATP), CDK11B/cyclin K (5 μM ATP), CDK12 (R722C)/cyclin K (5 μM ATP), CDK12/cyclin K (30 μM ATP), CDK13/cyclin K (5 μM ATP), CDK14/cyclin Y (PFTK1) (15 μM ATP), CDK15/cyclin A2 (100 μM ATP), CDK15/cyclin B1 (50 μM ATP), CDK16/cyclin (PCTAIRE) (10 μM ATP), CDK17/cyclin Y (PCTK2) (20 μM ATP), CDK18/cyclin Y (PCTK3) (20 μM ATP), CDK19/cyclin C (15 μM ATP). The percent enzyme activity relative to DMSO controls (set to 100%) was reported for all CDK/cyclin pairs.

### BioID

BioID (proximity-dependent <u>bio</u>tin <u>id</u>entification) studies were performed using the method described in Sears et al. (75). Briefly, HEK293T cells were seeded in 10 cm dish at 2.2 × 10^6^ cells and after 24 hours were transfected with 10 μg of plasmids encoding TFE3 and variants cloned into the expression vector pLX_313 (Broad Institute) and tagged with the mutated form of biotin ligase BirA (R118G) (or BirA*). Two days post transfection, 50 μM of Biotin (SigmaAldrich, #58855) was added to the plates. After 16-18 hours of biotin labeling, the cells were washed 2X with PBS and lysed using lysis buffer (50mM Tris-HCl, pH 7.5, 8M urea, 1 mM DTT, 1X Halt^TM^ protease inhibitor cocktail (ThermoFisher, #87786)). Pierce^TM^ Universal Nuclease (ThermoFisher, #88700) was added, mixed and incubated for 10 minutes at room temperature followed by addition of 1% Triton-X100 (ChemCruz, #sc-29112A). All subsequent steps were performed at 4°C. First, the lysate was sonicated (30 seconds for 3 rounds at 30% duty cycle, output level 3 using Branson Sonifier-250 tip sonicator) with one minute of ice incubation in between sonication to prevent heating. After the 3^rd^ sonication, lysis buffer (2X the lysate volume) was added and the lysate was sonicated for 1 minute followed by centrifugation at 16,600g for 10 minutes at 4°C. The lysate was pre-cleared using gelatin-sepharose beads (GE Healthcare, 17095601) for 2 hours at 4°C before binding biotinylated proteins using streptavidin-conjugated sepharose beads (GE Healthcare, 17511301) for 4 hours at 4°C. The beads were then washed 4 times with wash buffer (8M urea in 50 mM Tris-HCl) and once with 50 mM ammonium bicarbonate buffer and finally resuspended in 100 μL of 50mM ammonium bicarbonate with 1 mM Biotin and sent for mass spectrometry (Taplin Mass Spectrometry Facility, Harvard Medical School).

### Homology model of the DNA-bound TFE3 dimer

The published dimeric structure of TFE3 (PDB ID: 7F09) (14) does not have the DNA-binding region fully resolved. Hence, a homology model based on the homologous MITF protein bound to M-box (CATGTG) DNA (PDB ID: 7D8T) (45) was built to better understand the interactions of the TFE3 dimer with DNA. Chains A and B of the MITF structure were used as templates for the dimeric TFE3 homology model, using Prime (v. 7.4) as implemented in the Schrödinger Suite (Schrödinger, release 2023-4). The M-box DNA structures, found in Chains C and D of the template PDB, were transferred to make a combined homology model. Finally, the structure was subject to energy minimization using the protein preparation tool in the Schrödinger Software package.

## Acknowledgements

S.R.V.: acknowledges research support from Bayer U.S. LLC, Pharmaceuticals for this project in addition to support from the NCI R01CA286652; R01CA279044. P.Konda received funding from DoD KCRP Postdoctoral and Clinical Fellowship (HT94252310066). Experimental schematics were drawn using BioRender.com.

## Author contributions

Designed and performed experiments and analysis: D.R.T., R.D., P.Khanna., A.S., P.S., M.A., J.L., A.L.; Data analysis: P.Konda., Y.C., M.Z.; Structure analysis: J.B., C.P.; Designed, collaborated, and supervised aspects of the study: J.M.T., D.F., S.G., J.C., R.R., O.G.; Manuscript writing with input from all authors: D.R.T and S.R.V.; Designed and supervised the overall study: S.R.V.

## References

1. Ellati RT, Abukhiran I, Alqasem K, Jasser J, Khzouz J, Bisharat T, et al. Clinicopathologic Features of Translocation Renal Cell Carcinoma. Clin Genitourin Cancer. 2017 Feb;15(1):112–6.

2. Geller JI, Ehrlich PF, Cost NG, Khanna G, Mullen EA, Gratias EJ, et al. Characterization of adolescent and pediatric renal cell carcinoma: A report from the Children’s Oncology Group study AREN03B2. Cancer. 2015;121(14):2457–64.

3. Classe M, Malouf GG, Su X, Yao H, Thompson EJ, Doss DJ, et al. Incidence, clinicopathological features and fusion transcript landscape of translocation renal cell carcinomas. Histopathology. 2017 Jun;70(7):1089–97.

4. Meyer PN, Clark JI, Flanigan RC, Picken MM. Xp11.2 translocation renal cell carcinoma with very aggressive course in five adults. Am J Clin Pathol. 2007 Jul;128(1):70–9.

5. Bakouny Z, Sadagopan A, Ravi P, Metaferia NY, Li J, AbuHammad S, et al. Integrative clinical and molecular characterization of translocation renal cell carcinoma. Cell Rep. 2022 Jan 4;38(1):110190.

6. Achom M, Sadagopan A, Bao C, McBride F, Li J, Konda P, et al. A genetic basis for sex differences in Xp11 translocation renal cell carcinoma. Cell. 2024 Oct 3;187(20):5735–5752.e25.

7. Sun G, Chen J, Liang J, Yin X, Zhang M, Yao J, et al. Integrated exome and RNA sequencing of TFE3-translocation renal cell carcinoma. Nat Commun. 2021 Sep 6;12(1):5262.

8. Qu Y, Wu X, Anwaier A, Feng J, Xu W, Pei X, et al. Proteogenomic characterization of MiT family translocation renal cell carcinoma. Nat Commun. 2022 Dec 5;13(1):7494.

9. Kauffman EC, Ricketts CJ, Rais-Bahrami S, Yang Y, Merino M, Bottaro D, et al. Molecular Genetics and Cellular Characteristics of TFE3 and TFEB Translocation Renal Cell Carcinomas. Nat Rev Urol. 2014 Aug;11(8):465–75.

10. Bakouny Z, Sadagopan A, Ravi P, Metaferia NY, Li J, AbuHammad S, et al. Integrative clinical and molecular characterization of translocation renal cell carcinoma. Cell Rep. 2022 Jan 4;38(1):110190.

11. Malouf GG, Su X, Yao H, Gao J, Xiong L, He Q, et al. Next-generation sequencing of translocation renal cell carcinoma reveals novel RNA splicing partners and frequent mutations of chromatin-remodeling genes. Clin Cancer Res Off J Am Assoc Cancer Res. 2014 Aug 1;20(15):4129–40.

12. Beckmann H, Su LK, Kadesch T. TFE3: a helix-loop-helix protein that activates transcription through the immunoglobulin enhancer muE3 motif. Genes Dev. 1990 Feb 1;4(2):167–79.

13. Martina JA, Diab HI, Lishu L, Jeong-A L, Patange S, Raben N, et al. The Nutrient-Responsive Transcription Factor TFE3, Promotes Autophagy, Lysosomal Biogenesis, and Clearance of Cellular Debris. Sci Signal. 2014 Jan 21;7(309):ra9.

14. Yang G, Li P, Liu Z, Wu S, Zhuang C, Qiao H, et al. Structural basis for the dimerization mechanism of human transcription factor E3. Biochem Biophys Res Commun. 2021 Sep;569:41–6.

15. Roman C, Matera AG, Cooper C, Artandi S, Blain S, Ward DC, et al. mTFE3, an X-linked transcriptional activator containing basic helix-loop-helix and zipper domains, utilizes the zipper to stabilize both DNA binding and multimerization. Mol Cell Biol. 1992 Feb;12(2):817–27.

16. Simonaggio A, Ambrosetti D, Verkarre V, Auvray M, Oudard S, Vano YA. MiTF/TFE Translocation Renal Cell Carcinomas: From Clinical Entities to Molecular Insights. Int J Mol Sci. 2022 Jul 11;23(14):7649.

17. Bushweller JH. Targeting transcription factors in cancer — from undruggable to reality. Nat Rev Cancer. 2019 Nov;19(11):611–24.

18. Henley MJ, Koehler AN. Advances in targeting “undruggable” transcription factors with small molecules. Nat Rev Drug Discov. 2021 Sep;20(9):669–88.

19. Asante Y, Benischke K, Osman I, Ngo QA, Wurth J, Laubscher D, et al. PAX3-FOXO1 uses its activation domain to recruit CBP/P300 and shape RNA Pol2 cluster distribution. Nat Commun. 2023 Dec 15;14(1):8361.

20. Olsen SN, Godfrey L, Healy JP, Choi YA, Kai Y, Hatton C, et al. MLL::AF9 degradation induces rapid changes in transcriptional elongation and subsequent loss of an active chromatin landscape. Mol Cell. 2022 Mar 17;82(6):1140–1155.e11.

21. Richters A, Doyle SK, Freeman DB, Lee C, Leifer BS, Jagannathan S, et al. Modulating Androgen Receptor-Driven Transcription in Prostate Cancer with Selective CDK9 Inhibitors. Cell Chem Biol. 2021 Feb;28(2):134–147.e14.

22. Bradner JE, Hnisz D, Young RA. Transcriptional Addiction in Cancer. Cell. 2017 Feb;168(4):629–43.

23. Wyce A, Degenhardt Y, Bai Y, Le B, Korenchuk S, Crouthamel MC, et al. Inhibition of BET bromodomain proteins as a therapeutic approach in prostate cancer. Oncotarget. 2013 Dec 31;4(12):2419–29.

24. Raina K, Lu J, Qian Y, Altieri M, Gordon D, Rossi AMK, et al. PROTAC-induced BET protein degradation as a therapy for castration-resistant prostate cancer. Proc Natl Acad Sci. 2016 Jun 28;113(26):7124–9.

25. Dawson MA, Prinjha RK, Dittmann A, Giotopoulos G, Bantscheff M, Chan WI, et al. Inhibition of BET recruitment to chromatin as an effective treatment for MLL-fusion leukaemia. Nature. 2011 Oct;478(7370):529–33.

26. Zuber J, Shi J, Wang E, Rappaport AR, Herrmann H, Sison EA, et al. RNAi screen identifies Brd4 as a therapeutic target in acute myeloid leukaemia. Nature. 2011 Oct;478(7370):524–8.

27. Delmore JE, Issa GC, Lemieux ME, Rahl PB, Shi J, Jacobs HM, et al. BET Bromodomain Inhibition as a Therapeutic Strategy to Target c-Myc. Cell. 2011 Sep;146(6):904–17.

28. Grembecka J, He S, Shi A, Purohit T, Muntean AG, Sorenson RJ, et al. Menin-MLL Inhibitors Reverse Oncogenic Activity of MLL Fusion Proteins in Leukemia. Nat Chem Biol. 2012 Jan 29;8(3):277–84.

29. Daigle SR, Olhava EJ, Therkelsen CA, Basavapathruni A, Jin L, Boriack-Sjodin PA, et al. Potent inhibition of DOT1L as treatment of MLL-fusion leukemia. Blood. 2013 Aug 8;122(6):1017–25.

30. Garber K. Menin inhibitors seek to debut as newest targeted therapy for leukaemia. Nat Rev Drug Discov. 2024 Aug;23(8):567–9.

31. Wang Y, Zhang T, Kwiatkowski N, Abraham BJ, Lee TI, Xie S, et al. CDK7-Dependent Transcriptional Addiction in Triple-Negative Breast Cancer. Cell. 2015 Sep 24;163(1):174–86.

32. Toure MA, Motoyama K, Xiang Y, Urgiles J, Kabinger F, Koglin AS, et al. Targeted degradation of CDK9 potently disrupts the MYC-regulated network. Cell Chem Biol. 2025 Apr;32(4):542–555.e10.

33. Huang CH, Lujambio A, Zuber J, Tschaharganeh DF, Doran MG, Evans MJ, et al. CDK9-mediated transcription elongation is required for MYC addiction in hepatocellular carcinoma.

34. Picco G, Cattaneo CM, Van Vliet EJ, Crisafulli G, Rospo G, Consonni S, et al. Werner Helicase Is a Synthetic-Lethal Vulnerability in Mismatch Repair–Deficient Colorectal Cancer Refractory to Targeted Therapies, Chemotherapy, and Immunotherapy. Cancer Discov. 2021 Aug 1;11(8):1923–37.

35. Chan EM, Shibue T, McFarland JM, Gaeta B, Ghandi M, Dumont N, et al. WRN helicase is a synthetic lethal target in microsatellite unstable cancers. Nature. 2019 Apr;568(7753):551–6.

36. Rodríguez Pérez F, Natwick D, Schiff L, McSwiggen D, Heckert A, Huey M, et al. WRN inhibition leads to its chromatin-associated degradation via the PIAS4-RNF4-p97/VCP axis. Nat Commun. 2024 Jul 18;15:6059.

37. Guan J, Zhou W, Hafner M, Blake RA, Chalouni C, Chen IP, et al. Therapeutic Ligands Antagonize Estrogen Receptor Function by Impairing Its Mobility. Cell. 2019 Aug;178(4):949–963.e18.

38. Zhan Y, Kost-Alimova M, Shi X, Leo E, Bardenhagen JP, Shepard HE, et al. Development of novel cellular histone-binding and chromatin-displacement assays for bromodomain drug discovery. Epigenetics Chromatin. 2015;8:37.

39. Beckmann H, Kadesch T. The leucine zipper of TFE3 dictates helix-loop-helix dimerization specificity. Genes Dev. 1991 Jun;5(6):1057–66.

40. Steingrímsson E, Tessarollo L, Pathak B, Hou L, Arnheiter H, Copeland NG, et al. Mitf and Tfe3, two members of the Mitf-Tfe family of bHLH-Zip transcription factors, have important but functionally redundant roles in osteoclast development. Proc Natl Acad Sci. 2002 Apr 2;99(7):4477–82.

41. Tsherniak A, Vazquez F, Montgomery PG, Weir BA, Kryukov G, Cowley GS, et al. Defining a Cancer Dependency Map. Cell. 2017 Jul 27;170(3):564–576.e16.

42. DepMap: The Cancer Dependency Map Project at Broad Institute [Internet]. [cited 2025 Jun 2]. Available from: https://depmap.org/portal/

43. Dempster JM, Boyle I, Vazquez F, Root DE, Boehm JS, Hahn WC, et al. Chronos: a cell population dynamics model of CRISPR experiments that improves inference of gene fitness effects. Genome Biol. 2021 Dec 20;22(1):343.

44. Li B, Sadagopan A, Li J, Wu Y, Cui Y, Konda P, et al. A framework for target discovery in rare cancers. BioRxiv Prepr Serv Biol. 2024 Nov 20;2024.10.24.620074.

45. Liu Z, Chen K, Dai J, Xu P, Sun W, Liu W, et al. A unique hyperdynamic dimer interface permits small molecule perturbation of the melanoma oncoprotein MITF for melanoma therapy. Cell Res. 2023 Jan;33(1):55–70.

46. Filippakopoulos P, Qi J, Picaud S, Shen Y, Smith WB, Fedorov O, et al. Selective inhibition of BET bromodomains. Nature. 2010 Dec 23;468(7327):1067–73.

47. Nardone C, Palanski BA, Scott DC, Timms RT, Barber KW, Gu X, et al. A central role for regulated protein stability in the control of TFE3 and MITF by nutrients. Mol Cell. 2023 Jan;83(1):57–73.e9.

48. Li B, Sadagopan A, Li J, Wu Y, Cui Y, Konda P, et al. A framework for target discovery in rare cancers [Internet]. 2024 [cited 2025 Jan 24]. Available from: http://biorxiv.org/lookup/doi/10.1101/2024.10.24.620074

49. Kanda H, Yamawaki K. Bardoxolone methyl: drug development for diabetic kidney disease. Clin Exp Nephrol. 2020 Oct;24(10):857–64.

50. Wang S, Griffiths G, Midgley CA, Barnett AL, Cooper M, Grabarek J, et al. Discovery and Characterization of 2-Anilino-4-(Thiazol-5-yl)Pyrimidine Transcriptional CDK Inhibitors as Anticancer Agents. Chem Biol. 2010 Oct;17(10):1111–21.

51. Cidado J, Boiko S, Proia T, Ferguson D, Criscione SW, San Martin M, et al. AZD4573 Is a Highly Selective CDK9 Inhibitor That Suppresses MCL-1 and Induces Apoptosis in Hematologic Cancer Cells. Clin Cancer Res. 2020 Feb 15;26(4):922–34.

52. Malumbres M, Harlow E, Hunt T, Hunter T, Lahti JM, Manning G, et al. Cyclin-dependent kinases: a family portrait. Nat Cell Biol. 2009 Nov;11(11):1275–6.

53. Baba M, Furuya M, Motoshima T, Lang M, Funasaki S, Ma W, et al. TFE3 Xp11.2 Translocation Renal Cell Carcinoma Mouse Model Reveals Novel Therapeutic Targets and Identifies GPNMB as a Diagnostic Marker for Human Disease. Mol Cancer Res. 2019 Aug;17(8):1613–26.

54. Richters A, Doyle SK, Freeman DB, Lee C, Leifer BS, Jagannathan S, et al. Modulating Androgen Receptor-Driven Transcription in Prostate Cancer with Selective CDK9 Inhibitors. Cell Chem Biol. 2021 Feb;28(2):134–147.e14.

55. Toure MA, Motoyama K, Xiang Y, Urgiles J, Kabinger F, Koglin AS, et al. Targeted Degradation of CDK9 Potently Disrupts the MYC Transcriptional Network [Internet]. bioRxiv; 2024 [cited 2025 Feb 26]. p. 2024.05.14.593352. Available from: https://www.biorxiv.org/content/10.1101/2024.05.14.593352v1

56. Diamond JR, Boni V, Lim E, Nowakowski G, Cordoba R, Morillo D, et al. First-in-Human Dose-Escalation Study of Cyclin-Dependent Kinase 9 Inhibitor VIP152 in Patients with Advanced Malignancies Shows Early Signs of Clinical Efficacy. Clin Cancer Res. 2022 Apr 1;28(7):1285–93.

57. Frigault MM, Mithal A, Wong H, Stelte-Ludwig B, Mandava V, Huang X, et al. Enitociclib, a Selective CDK9 Inhibitor, Induces Complete Regression of MYC+ Lymphoma by Downregulation of RNA Polymerase II Mediated Transcription. Cancer Res Commun. 2023 Nov 9;3(11):2268–79.

58. Choueiri TK, Lim ZD, Hirsch MS, Tamboli P, Jonasch E, McDermott DF, et al. Vascular endothelial growth factor-targeted therapy for the treatment of adult metastatic Xp11.2 translocation renal cell carcinoma. Cancer. 2010;116(22):5219–25.

59. Malouf GG, Camparo P, Oudard S, Schleiermacher G, Theodore C, Rustine A, et al. Targeted agents in metastatic Xp11 translocation/TFE3 gene fusion renal cell carcinoma (RCC): a report from the Juvenile RCC Network. Ann Oncol. 2010 Sep;21(9):1834–8.

60. Alhalabi O, Thouvenin J, Négrier S, Vano YA, Campedel L, Hasanov E, et al. Immune Checkpoint Therapy Combinations in Adult Advanced MiT Family Translocation Renal Cell Carcinomas. The Oncologist. 2023 May 8;28(5):433–9.

61. Peterlin BM, Price DH. Controlling the Elongation Phase of Transcription with P-TEFb. Mol Cell. 2006 Aug;23(3):297–305.

62. Jonkers I, Lis JT. Getting up to speed with transcription elongation by RNA polymerase II. Nat Rev Mol Cell Biol. 2015 Mar;16(3):167–77.

63. Sonawane YA, Taylor MA, Napoleon JV, Rana S, Contreras JI, Natarajan A. Cyclin Dependent Kinase 9 Inhibitors for Cancer Therapy. J Med Chem. 2016 Oct 13;59(19):8667–84.

64. Wang S, Griffiths G, Midgley CA, Barnett AL, Cooper M, Grabarek J, et al. Discovery and Characterization of 2-Anilino-4-(Thiazol-5-yl)Pyrimidine Transcriptional CDK Inhibitors as Anticancer Agents. Chem Biol. 2010 Oct;17(10):1111–21.

65. Byrd JC, Lin TS, Dalton JT, Wu D, Phelps MA, Fischer B, et al. Flavopiridol administered using a pharmacologically derived schedule is associated with marked clinical efficacy in refractory, genetically high-risk chronic lymphocytic leukemia. Blood. 2007 Jan 15;109(2):399–404.

66. Frigault MM, Mithal A, Wong H, Stelte-Ludwig B, Mandava V, Huang X, et al. Enitociclib, a Selective CDK9 Inhibitor, Induces Complete Regression of MYC+ Lymphoma by Downregulation of RNA Polymerase II Mediated Transcription. Cancer Res Commun. 2023 Nov 9;3(11):2268–79.

67. Tran S, Sipila P, Frigault MM, Stelte-Ludwig B, Johnson AJ, Birkett J, et al. Enitociclib, a selective CDK9 inhibitor: in vitro and in vivo preclinical studies in multiple myeloma. Blood Neoplasia. 2025 Feb;2(1):100050.

68. Mandal R, Becker S, Strebhardt K. Targeting CDK9 for Anti-Cancer Therapeutics. Cancers. 2021 May 1;13(9):2181.

69. Onji H, Murai J. Reconsidering the mechanisms of action of PARP inhibitors based on clinical outcomes. Cancer Sci. 2022 Sep;113(9):2943–51.

70. Kaelin WG. Common pitfalls in preclinical cancer target validation. Nat Rev Cancer. 2017 Jul;17(7):441–50.

71. DepMap, Broad (2024). Current DepMap Release data, including CRISPR Screens, PRISM Drug Screens, Copy Number, Mutation, Expression, and Fusions. DepMap 23Q2 Public. Figshare+. Dataset.;

72. Huan C, Sashital D, Hailemariam T, Kelly ML, Roman CAJ. Renal Carcinoma-associated Transcription Factors TFE3 and TFEB Are Leukemia Inhibitory Factor-responsive Transcription Activators of E-cadherin. J Biol Chem. 2005 Aug;280(34):30225–35.

73. Viswanathan SR, Nogueira MF, Buss CG, Krill-Burger JM, Wawer MJ, Malolepsza E, et al. Genome-scale analysis identifies paralog lethality as a vulnerability of chromosome 1p loss in cancer. Nat Genet. 2018 Jul;50(7):937–43.

74. Ritchie ME, Phipson B, Wu D, Hu Y, Law CW, Shi W, et al. limma powers differential expression analyses for RNA-sequencing and microarray studies. Nucleic Acids Res. 2015 Apr 20;43(7):e47–e47.

75. Sears RM, May DG, Roux KJ. BioID as a Tool for Protein-Proximity Labeling in Living Cells. Methods Mol Biol Clifton NJ. 2019;2012:299–313.

