## Supplemental Figures for "Phenotypic screening converges on CDK9 inhibition as a therapeutic strategy in translocation renal cell carcinoma"

### Supplementary Figures and Captions

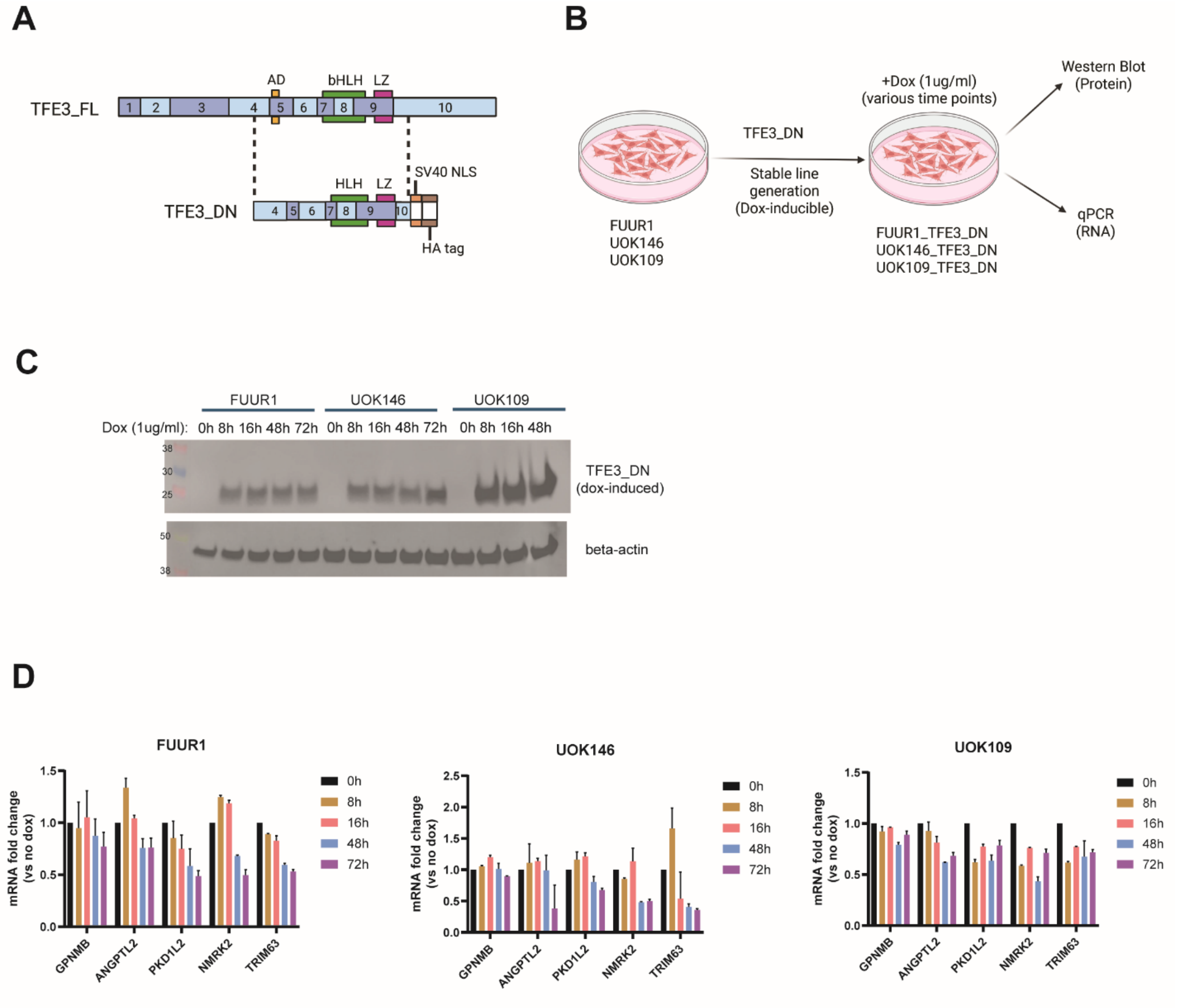


Figure S1

### Supplementary Figure S1. Expression of TFE3 dominant negative (TFE3_DN) suppress TFE3 fusion activity in tRCC.

**(A)** Schematic of TFE3_DN construct (72).

**(B)** Experiment outline for dox-inducible TFE3_DN tRCC stable line generation and for protein (Western blot) and mRNA (qPCR) studies.

**(C)** Validation of doxycycline-inducible TFE3_DN expression by Western blot in three tRCC lines (FUUR1, *ASPSCR1-TFE3*; UOK146, *PRCC-TFE3*; UOK109, *NONO-TFE3*).

**(D)** qPCR analysis of TFE3 target genes mRNA level after TFE3_DN expression in tRCC lines (mean ± SEM, n=2).

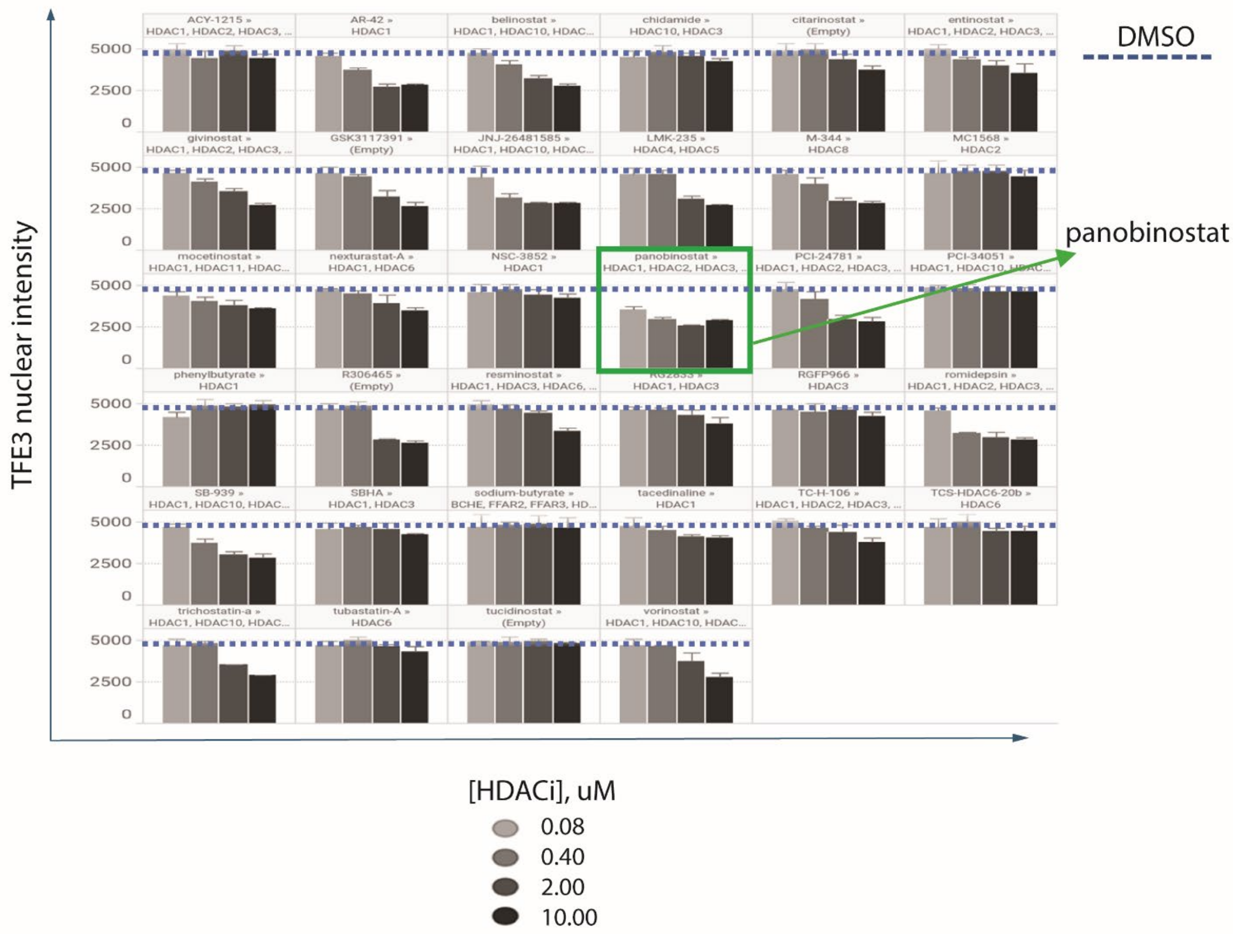
DMSO
panobinostat
TFE3 nuclear intensity
5000
2500
0
ACY-1215 » HDAC1, HDAC2, HDAC3, ...
AR-42 » HDAC1
belinostat » HDAC1, HDAC10, HDAC...
chidamide » HDAC10, HDAC3
citarinostat » (Empty)
entinostat » HDAC1, HDAC2, HDAC3, ...
givinostat » HDAC1, HDAC2, HDAC3, ...
GSK3117391 » (Empty)
JNJ-26481585 » HDAC1, HDAC10, HDAC...
LMK-235 » HDAC4, HDAC5
M-344 » HDAC8
MC1568 » HDAC2
mocetinostat » HDAC1, HDAC11, HDAC...
nexturastat-A » HDAC1, HDAC6
NSC-3852 » HDAC1
panobinostat » HDAC1, HDAC2, HDAC3, ...
PCI-24781 » HDAC1, HDAC2, HDAC3, ...
PCI-34051 »
phenylbutyrate » HDAC1
R306465 » (Empty)
resminostat » HDAC1, HDAC3, HDAC6, ...
HDAC1, HDAC3
RGFP966 » HDAC3
romidepsin » HDAC1, HDAC2, HDAC3, ...
SB-939 » HDAC1, HDAC10, HDAC...
SBHA » HDAC1, HDAC3
sodium-butyrate » BCHE, FFAR2, FFAR3, HD...
tacedinaline » HDAC1
TC-H-106 » HDAC1, HDAC2, HDAC3, ...
TCS-HDAC6-20b » HDAC6
trichostatin-a » HDAC1, HDAC10, HDAC...
tubastatin-A » HDAC6
tucidinostat » (Empty)
vorinostat » HDAC1, HDAC10, HDAC...
[HDACi], uM
0.08
0.40
2.00
10.00

**Figure S2**

**Supplementary Figure S2. TFE3 chromatin displacement by HDAC inhibitors identifies panobinostat as TFE3 displacement control.**

Dose response for TFE3 nuclear displacement by various HDAC inhibitors included in the EpiMod library. Panobinostat (green box) shows the strongest displacement with activity even at the lowest compound concentration of 0.08 μM.

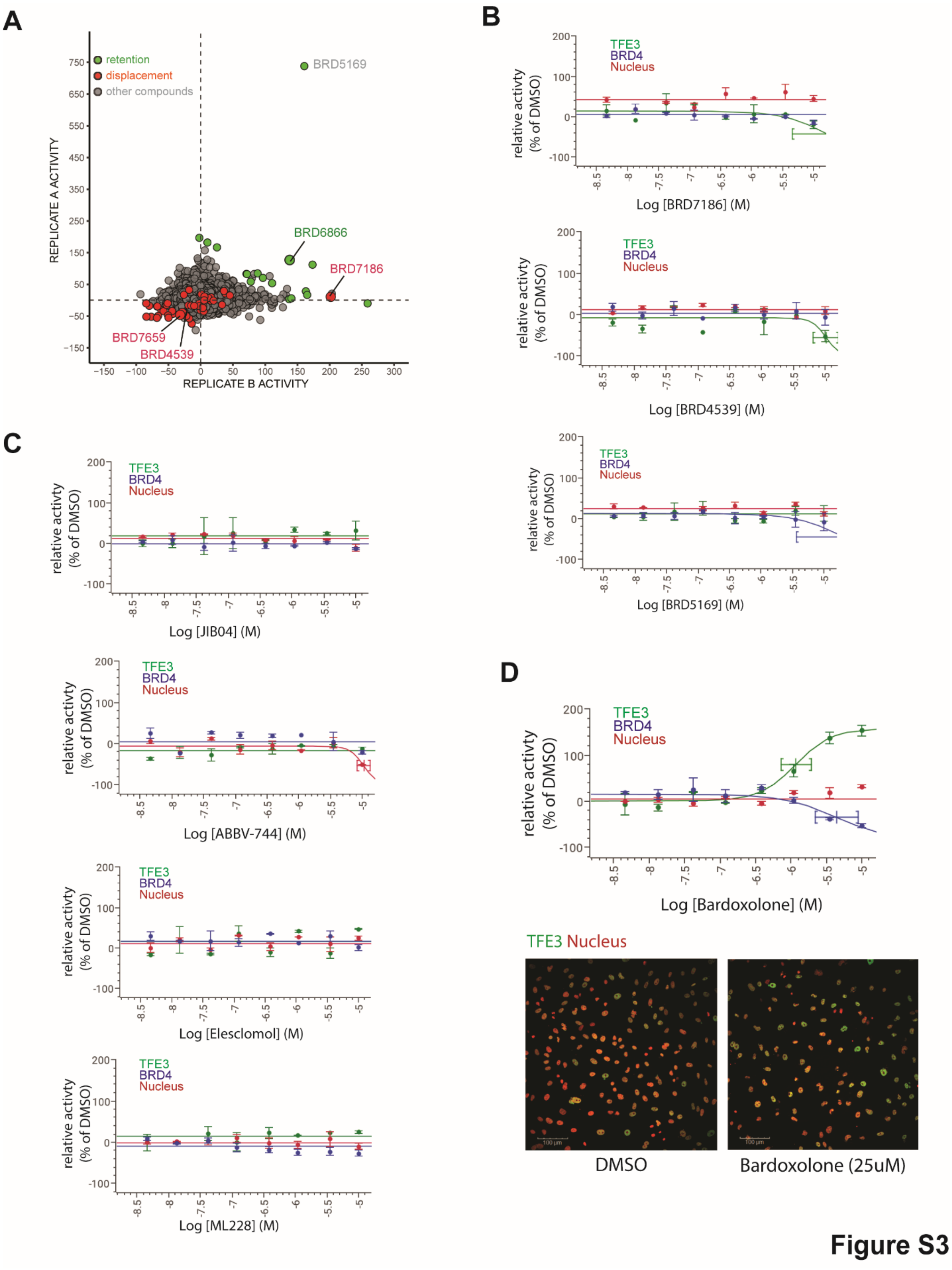
A
retention
displacement
other compounds
BRD5169
BRD6866
BRD7186
BRD7659
BRD4539
REPLICATE A ACTIVITY
REPLICATE B ACTIVITY
B
TFE3
BRD4
Nucleus
relative activty (% of DMSO)
Log [BRD7186] (M)
Log [BRD4539] (M)
Log [BRD5169] (M)
C
Log [JIB04] (M)
Log [ABBV-744] (M)
Log [Elesclomol] (M)
Log [ML228] (M)
D
Log [Bardoxolone] (M)
TFE3 Nucleus
DMSO
Bardoxolone (25uM)

**Figure S3**

### Supplementary Figure S3. Validation of hits using TFE3 chromatin displacement assay.

**(A)** Replicate-Replicate scatterplot for all 5 runs combined for chromatin displacement screening of 25,000 compounds in in FUUR1 cells. Chromatin displacers, red; chromatin retention, green (see methods for hit selection that was performed individually from each run). Cumulative primary hits are labeled in red or green; validated hits are named in green or red. Outlier hit is named in gray.

**(B)** Dose response curve in chromatin displacement assay from the displacement hits (BRD7186, BRD4539) and outlier hit (BRD5169) from the 25,000 compounds primary screen.

**(C)** Select hits from the repurposing library viability screen that showed tRCC-selective cell killing (48), and which were tested for TFE3 displacement using chromatin displacement assay.

**(D)** Bardoxolone, a hit from the repurposing library viability screen, shows a dose-dependent TFE3 retention phenotype in chromatin displacement assay (images show signal at DMSO vs. 25 $\mu$M).

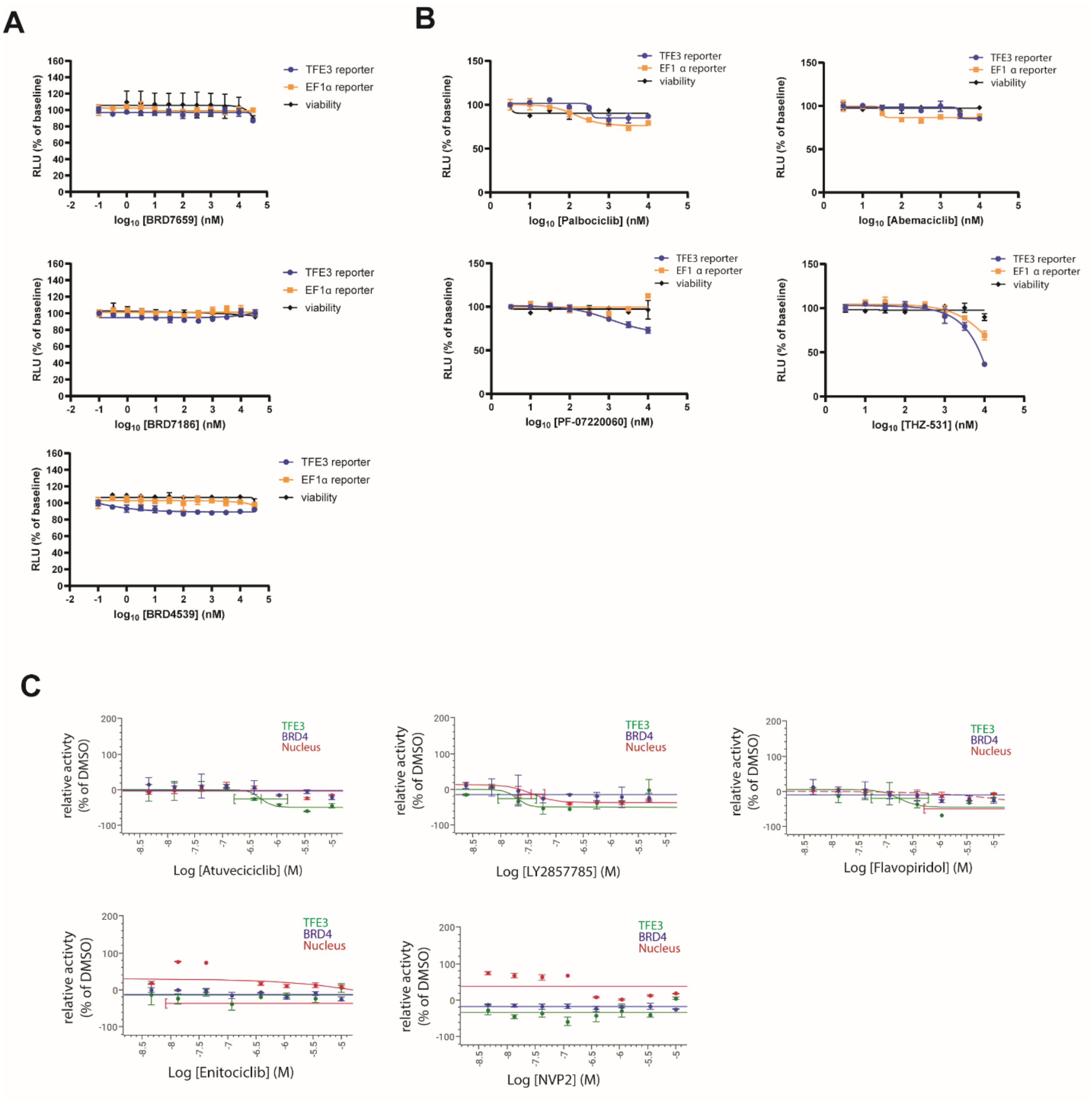
A
B
C
TFE3 reporter
EF1α reporter
viability
RLU (% of baseline)
log10 [BRD7659] (nM)
log10 [BRD7186] (nM)
log10 [BRD4539] (nM)
log10 [Palbociclib] (nM)
log10 [Abemaciclib] (nM)
log10 [PF-07220060] (nM)
log10 [THZ-531] (nM)
TFE3
BRD4
Nucleus
relative activty (% of DMSO)
Log [Atuveciciclib] (M)
Log [LY2857785] (M)
Log [Flavopiridol] (M)
Log [Enitociclib] (M)
Log [NVP2] (M)

**Figure S4**

#### Supplementary Figure S4. TFE3 chromatin displacement by CDK9 inhibitors.

**(A)** Dual reporter luciferase assay in UOK109 cells with BRD7659, BRD7186, and BRD4539 (mean ± SD, n=3).

**(B)** Dual reporter luciferase assay in UOK109 cells with a CDK4/6 and CDK12/13 inhibitors (mean ± SD, n=2).

**(C)** Dose-response of various CDK9 inhibitors in the chromatin displacement assay. Data and image are color coded for TFE3 (green), BRD4 (blue), nucleus (red) (mean ± SD, n=3).

### Supplementary Table Captions

**Supplementary Table S1.** Screening results from the EpiMod library showing mean of nuclear TFE3 (AF-488 signal) intensity and nuclei count for all compounds tested at four different concentrations (10μM, 2μM, 0.4μM, and 0.08μM).

**Supplementary Table S2.** List of compounds that were validated in TFE3 chromatin displacement assay using a 8-point compound dose range. The list includes the primary hits from the 25,000 library CDA screening study (n=62), select hits from the repurposing library screening study (48) showing selective viability effect on tRCC compared to ccRCC lines (n=5), and select CDK 7/9 inhibitors (n=6).
